# Tissue tropisms of avian influenza A viruses affect their spillovers from wild birds to pigs

**DOI:** 10.1101/2020.04.14.040964

**Authors:** Xiaojian Zhang, Fred L. Cunningham, Lei Li, Katie Hanson-Dorr, Liyuan Liu, Kaitlyn Waters, Minhui Guan, Alicia K. Olivier, Brandon S. Schmit, Jacqueline M. Nolting, Andrew S. Bowman, Mia Kim Torchetti, Thomas J. DeLiberto, Xiu-Feng Wan

**Author notes:** These authors contributed equally to this work (Xiaojian Zhang, Fred L. Cunningham). Correspondence: Dr. Thomas J. DeLiberto by or Dr. Xiu-Feng Wan by.

## Abstract

Wild aquatic birds maintain a large genetically diverse pool of influenza A viruses (IAVs), which can be transmitted to lower mammals and ultimately humans. Through phenotypic analyses, only a small set of avian IAVs replicated well in the epithelial cells of swine upper respiratory tracts, and these viruses were shown to infect and cause virus shedding in pigs. Such a phenotypic trait appears to emerge randomly and are distributed among IAVs across multiple avian species, geographic and temporal orders, and is determined not by receptor binding preference but other markers across genomic segments, such as those in the ribonucleoprotein complex. This study demonstrates that phenotypic variants exist among avian IAVs, only a few of which may result in viral shedding in pigs upon infection, providing opportunities for these viruses to become pig adapted, thus posing a higher potential risk for creating novel variants or detrimental reassortants within pig populations.

**Author Summary:** Having both avian-like receptors and human-like α2,6-linked sialic acid receptors, swine serve as a “mixing vessel” for generating human influenza pandemic strains. All HA subtypes of IAVs can infect swine; however, only sporadic cases of avian IAVs are reported in domestic swine. The molecular mechanisms affecting avian IAVs ability to infect swine are still not fully understood. Through phenotypic analyses, this study suggested that tissue tropisms (i.e., in swine upper respiratory tracts) of avian IAVs affect their spillovers from wild birds to pigs, and this phenotype was determined not by receptor binding preference but by other markers across genomic segments, such as those in the ribonucleoprotein complex. In addition, our results showed that such a phenotypic trait was sporadically and randomly distributed among IAVs across multiple avian species, geographic and temporal orders. This study suggested an efficient way for risk assessment of avian IAVs, such as in evaluating their potentials to be transmitted from avian to pigs.

## Introduction

Influenza A viruses (IAVs), a member in the *Orthomyxoviridae* family, are a negative single stranded RNA virus with eight gene segments, encoding at least 11 proteins. The nomenclatures of IAVs are determined based on antigenic properties of two surface glycoproteins, Hemagglutinin (HA) and Neuraminidase (NA). To date, 18 HA subtypes and 11 NA subtypes have been identified [1, 2]. Besides humans, pigs, dogs, and horses, IAVs have been recovered from variety of bird species, including at least 105 wild bird species of 26 different families [3]. Among wild birds, birds of wetlands and aquatic environments such as the *Anseriformes* (particularly ducks, geese, and swans) and *Charadriiformes* (particularly gulls, terns, and waders) constitute the major natural IAV reservoir [4]. They maintain a large IAV genetic pool, which contributes to the appearance of new IAVs in other birds, lower mammals, and ultimately humans. To date, 16 HA subtypes (H1-H16) and 9 NA subtypes (N1-N9) of IAVs have been identified in these wild aquatic birds [4–6].

Similar to avian species, swine are one of the most important natural hosts for IAVs. To date, only three predominant subtypes of IAVs (i.e., H1N1, H1N2 and H3N2) are enzootic in pigs [7, 8]. Having both avian-like receptors (α2,3-linked sialic acid, SA2,3Gal) and human-like receptors (α2,6-linked sialic acid, SA2,6Gal), swine may serve as an intermediate host “mixing vessels” for generating human influenza pandemic strains [9]. For example, the 2009 H1N1 pandemic IAV is a reassortant with HA, NP and NS from classical swine (North American) lineage, PB2 and PA from avian (North American) lineage, PB1 from human seasonal H3N2, and NA and M from Eurasian swine lineage [10]. Although these pandemic viruses have avian-origin genetic segments, how and when these avian genes are introduced into swine is not clear. Under laboratory conditions, all HA subtypes of IAVs can infect swine [11]. However, only sporadic cases of avian IAVs are detected in pigs, including subtype H1, H3, H4, H5, H6, H7 and H9, and most of these spillovers are transient [12–14]. Thus, as a key component of influenza pandemic preparedness, it will be critical to understand, among a large IAV genetic pool maintained by wild aquatic birds, which avian IAVs can be potentially transmitted to and subsequently spread in pigs.

Prior studies have demonstrated that receptor binding preference is one of the key factors affecting transmissibility of avian IAVs in mammals [15]. Acquired mutations in the HA receptor binding sites can switch receptor binding properties of avian IAVs and enhance virus transmissibility in mammals. For example, four amino acid substitutions in the HA receptor binding sites and one in the polymerase complex protein basic polymerase 2 (PB2), enabled H5N1 avian IAV to transmit in ferrets through airborne droplets [16]. Mutations at HA residues 222 (alanine to valine) and 228 (glycine to serine) increased the receptor binding preference of H6N6 avian IAVs to SA2,6Gal [17]. In addition to glycan receptor binding preference, genetic constellation of RNP complex and adaptative mutations in RNP genes, including PB1, have been shown to affect the host and tissue tropisms of IAVs [18–21]. It has been suggested that avian RNP complex has defects in its replication ability in human cells, and mutations that occur across PB2, PB1, PA, and NP of RNP complex are demonstrated to facilitate adaptation of avian IAVs in humans and other mammalian species (reviewed in Mänz et al. [22]). Nevertheless, the prerequisite for an avian IAV to be transmitted in mammals, such as pigs, is that the virus should be able to cause infection and viral shedding thereby allowing further infection and allowing the virus opportunities to acquire adaptive mutations and/or genetic reassortment, which could further facilitate virus circulation in pigs.

The study aimed to understand the molecular mechanism(s) by which avian IAVs infect pigs and further to determine, among a large IAV genetic pool maintained by wild aquatic birds, which avian IAVs are more likely to be transmitted and further adapted to pigs. Our previous study suggested that tissue tropism of an IAV, especially the replication efficiency on the upper swine respiratory tract epithelial cells (e.g. swine nasal cells [SNE]), is an important factor selecting for a transmissible reassortant in pigs [23]. Here we hypothesize that a large diversity in replication phenotypes on SNE cells are present in avian IAVs but without evolutionary pressure, and that the viruses with a high replication ability on SNE cells would be more likely to be transmitted in pigs. To test these hypotheses, we focused on a subtype H4N6 avian IAVs, which have been detected twice in domestic swine populations in North America [24, 25].

## Results

### H4N6 avian viruses have a large extent of diversity in replication efficiency on respiratory tract epithelial cells

In 2015, A/swine/Missouri/A01727926/2015 (abbreviated as MO/15), was recovered from a sick pig with influenza-like clinical signs due to a spillover of H4N6 avian viruses from wild birds to domestic pigs [24]. Phylogenetic analyses showed that, similar to the isolate from a prior spillover, A/Swine/Ontario/01911/99(H4N6) (abbreviated as ON/99) [26], MO/15 is also an isolate of avian origin North America lineage. However, the HA gene of MO/15 belongs to the genetic lineage IV whereas that of ON/99 to genetic lineage I, demonstrating these viruses were from two independent spillover incidences (Fig. 1). We hypothesize that H4N6 avian IAVs circulating in wild birds possess a large range of replication efficiencies in mammalian respiratory tract epithelial cells, and only a small portion of avian IAVs have phenotypes similar to the spillover viruses detected in pigs (i.e. MO/15 and ON/99). To test this hypothesis, a total of 115 H4N6 isolates (Table S1) from wild birds in North America were selected for phenotype analyses, and these viruses were selected to represent the diversity in genomic sequences, avian host species, sampling times, and sampling locations to maximize the possibility in covering the phenotypic diversity of H4N6 viruses in wild birds.

**Fig. 1.**
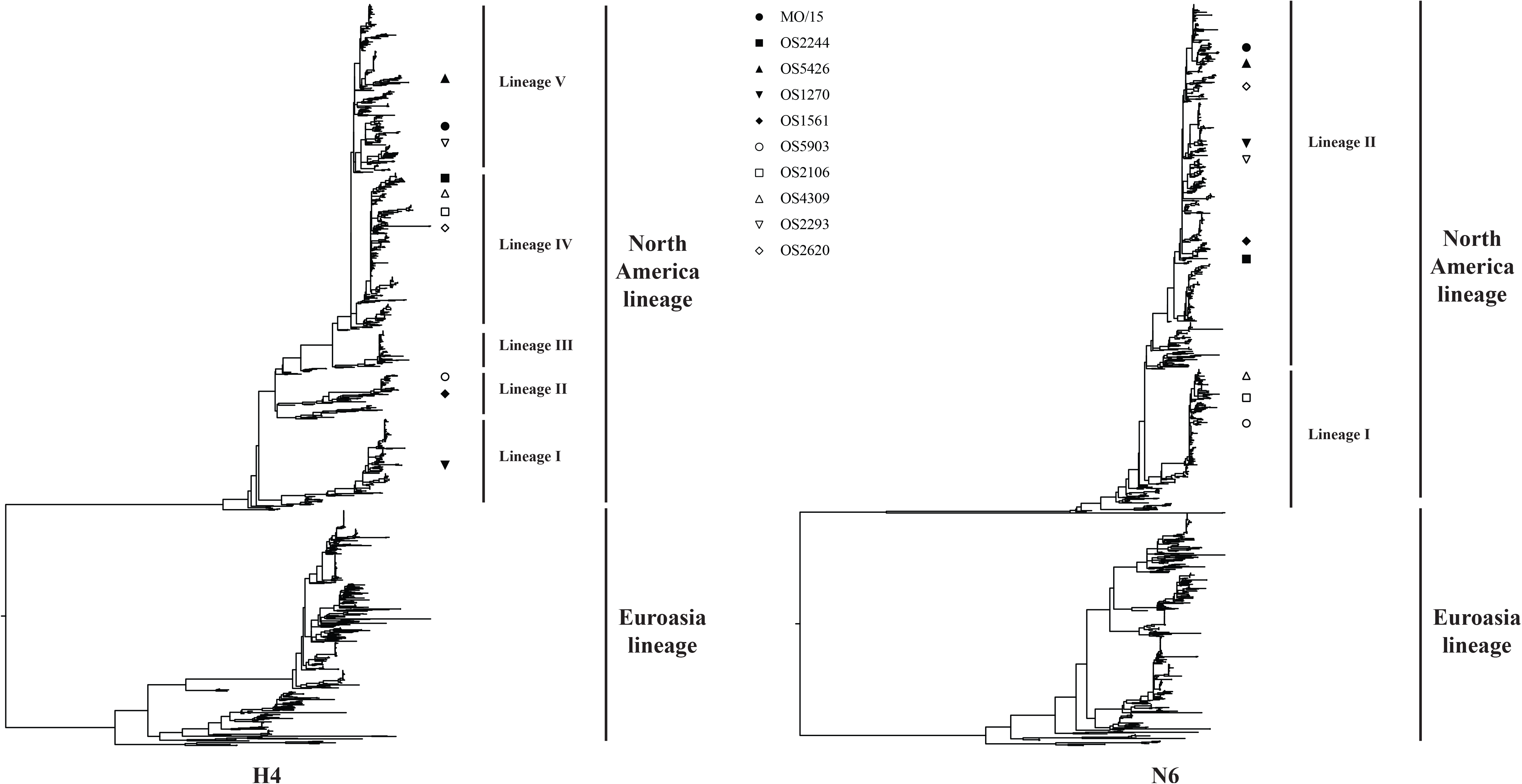
Phylogenetic analysis of HA and NA gene of H4N6 AIVs. The phylogenetic trees for each gene segment (HA and NA) was inferred using a maximum-likelihood method implemented in RAxML v8.2.9. A GAMMA model of rate heterogeneity and a generalized time-reversible (GTR) substitution model were applied in the analysis. Phylogenetic trees were then visualized by ggtree v1.6.11. The representative isolates were marked on the trees with symbols. Abbreviations: A/swine/Missouri/A01727926/2015 (H4N6), MO/15; A/blue-winged teal/Ohio/12OS2244/2012(H4N6), OS2244; A/blue-winged teal/Ohio/15OS5426/2015 (H4N6), OS5426; A/Mallard/Ohio/08OS1270/2008 (H4N6), OS1270; A/Blue winged teal/Illinois/10OS1561/2010 (H4N6), OS1561; A/American green winged teal/Mississippi/11OS5903/2011 (H4N6), OS5903; A/Blue winged teal/Ohio/12OS2106/2012 (H4N6), OS2106; A/Mallard/Wisconsin/11OS4309/2011 (H4N6), OS4309; /Blue winged teal/Ohio/12OS2293/2012, OS2293; and A/Blue winged teal/Missouri/11OS2620/2011 (H4N6), OS2620. Three isolates, MO/15, OS2244, and OS5426, were used in the animal study.

The replication phenotypic analyses were carried out on SNE, swine trachea epithelial cells (STE), and human A549 cells. The testing temperatures 33, 37, and 39 °C were used in SNE, STE, and A549 cells, respectively, to model the temperatures across the upper, middle, and lower respiratory tracts of the pig as previously performed [23]. Due to lack of available swine alveolar epithelial cells and the limited availability of swine lung primary cells, human A549 cells were used to model swine alveolar epithelial cells from swine lower respiratory tracks. Human A549 cells are characterized as a type II pulmonary epithelial cell to model IAV infection [27], and both human and swine type II pneumocytes express α-2,3- and α-2,6-linked sialic acid receptors [28, 29], and are targets of IAV infection [28, 30–32].

The 115 avian viruses showed a large diversity of replication efficiencies within and across the three cell models. These avian viruses replicated the best on A549 cells at 39 °C [5.875 (±standard deviation, ±1.426) Log_10_TCID_50_/mL], followed by STE cells at 37 °C [4.314 (±1.699) Log_10_TCID_50_/mL] and SNE cells at 33 °C [2.422 (±1.398) Log_10_TCID_50_/mL] (Fig. 2). There was a large diversity of the replication efficiencies across three cell model systems: using the SNE cells, only 43 testing viruses reached a viral titer of 2 Log_10_TCID_50_/mL or higher; with STE cells, 102 testing H4N6 viruses reached a viral titer of 2 Log_10_TCID_50_/mL or higher; and using A549 cells, 110 H4N6 IAVs reached a viral titer of 2 Log_10_TCID_50_/mL or higher.

**Fig. 2.**
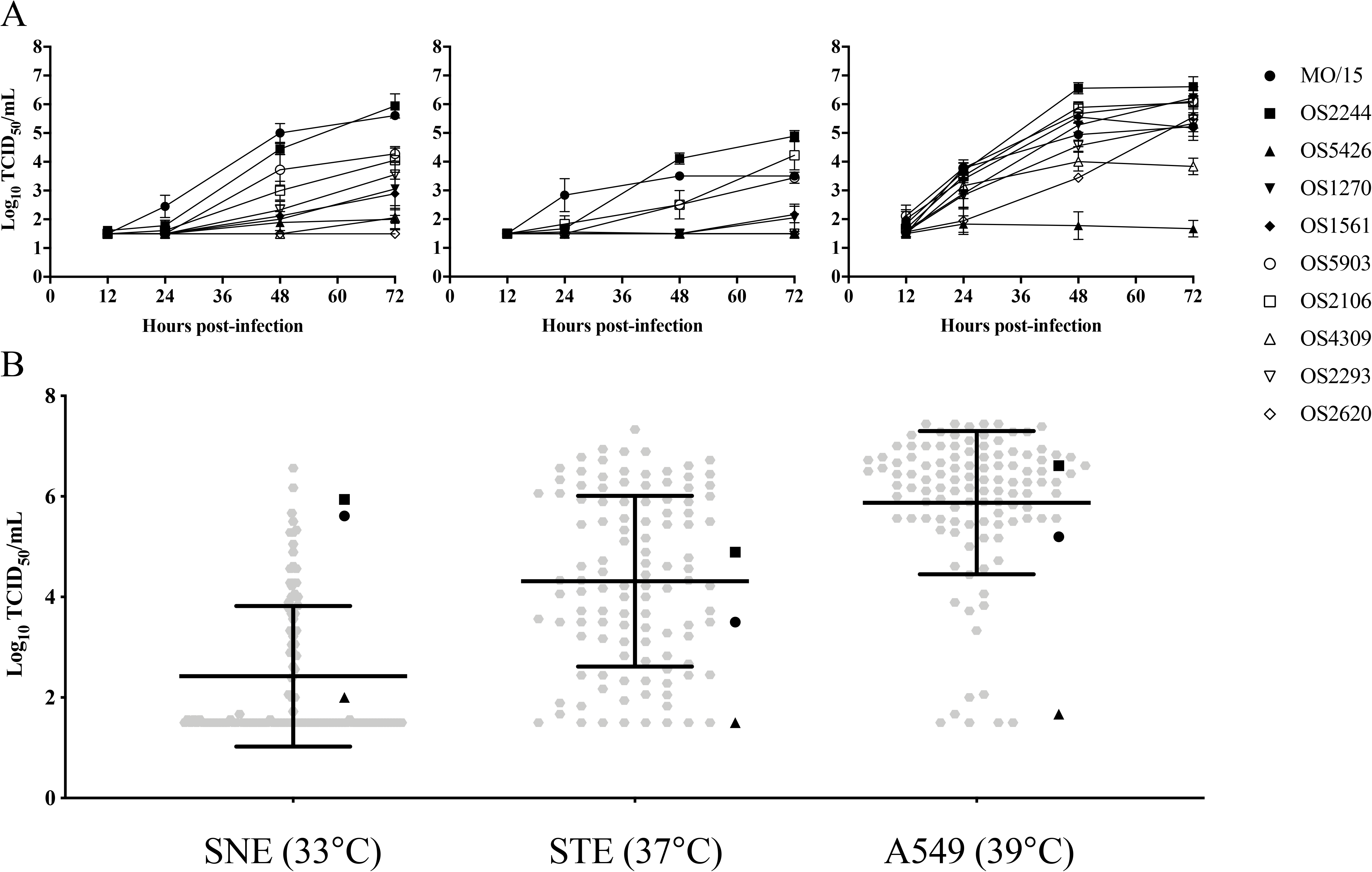
Growth dynamics of H4N6 IAVs in swine nasal epithelium cells (SNE), swine tracheal epithelium cells (STE), and human alveolar basal epithelial cells (A549). A) growth kinetics of nine representative H4N6 avian isolates and a H4N6 swine isolate, MO/15; B) summary of growth phenotypes on SNE, STE, and A549 cells at 72 h post-inoculation. Cells were infected at a multiplicity of infection of 0.001 TCID_50_/cell with the indicated viruses. Infected cells were incubated at 33°C, 37°C, or 39°C. Growth curves were determined by using the viral titers in the supernatants of infected cells obtained at 12, 24, 48, and 72 h post-inoculation. Data shown represent the mean titers ± standard errors (n = 3 cultures). Abbreviations are described in the legend of Fig. 1.

Results showed that the swine isolate MO/15 replicated efficiently in all three cell models, on SNE at 33 °C with a titer of 5.613 (±0.098) Log_10_TCID_50_/mL, on STE at 37 °C with a titer of 3.500 (±0.000) Log_10_TCID_50_/mL, and on A549 at 39 °C with a titer of 5.223 (±0.479) Log_10_TCID_50_/mL, respectively. Of the 115 testing viruses, only four had a growth titer greater than or equal to that of MO/15 at the same condition, especially in SNE. Twenty viruses replicated well across all three cells with a titer of ≥ 3.5 Log_10_TCID_50_/mL and showed a similar replication pattern to MO/15.

### Growth kinetics analyses support distinct replication patterns of H4N6 avian viruses on respiratory tract epithelial cells

To further differentiate phenotypic diversity of these viruses, we determined growth kinetics of MO/15 and nine other representative avian isolates that demonstrated distinct growth phenotypes in above analyses. In addition to the three cell models mentioned above, varied temperatures (i.e., 33, 37, and 39 °C) were to evaluate the effects of temperature on the growth of IAVs. Results demonstrated that there were significant variations among these 10 isolates in the growth kinetic patterns within and across the three cell models, which are consistent with those described above in the large-scale end-point replication efficiency analyses. Of interest, results showed that temperature had an effect on the viral growth kinetics in all three cell models. SNE cells supports virus replication at all three testing temperatures; STE cells supported virus replication at 37 °C better than the other two temperatures; and A549 cells supported high replication rates of H4N6 viruses at 37 and 39 °C but not 33 °C. In general, 33 °C, 37 °C, and 39 °C were shown to be optimal temperatures for virus replication at SNE, STE, and A549 cells, respectively (Figs. 2 and S1).

Results showed that MO/15 replicated efficiently in all three cell models (Fig. 2). Among the other nine avian viruses tested, A/blue-winged teal/Ohio/15OS5426/2015 (H4N6) (abbreviated as OS5426) replicated poorly in all three cell models, whereas A/blue-winged teal/Ohio/12OS2244/2012 (H4N6) (abbreviated as OS2244) replicated well in all, with peak viral titers of 5.943 (±0.418), 4.890 (±0.191), 6.610 (±0.348) Log_10_TCID_50_/mL at SNE (33 °C), STE (37 °C), and A549 (39 °C) cells, respectively.

Based on the phenotypic analyses, MO/15, OS5426 and OS2244 were selected for further phenotypic analysis of virus binding preference, polymerase activities, and infectivity and transmission in animals.

### The growth phenotypes of H4N6 avian viruses are affected by genetic diversity but genetic clade independent

To evaluate whether replication phenotypes of H4N6 viruses are associated with certain genetic clades of H4N6 viruses, we constructed phylogenetic trees of the selected H4N6 viruses and colored the viruses by their growth efficiency on the SNE cells at 33°C (Fig. 3). Results suggested, for each of the eight gene segments, the high growth strains are sporadically located across the phylogenetic tree, and there are no clear patterns identified between the viral growth efficiency and the genetic clades of any genes. The two spill-over strains, MO/15 and ON/99, are not genetically associated with high growth H4N6 avian strains identified in our phenotypic analyses. No patterns were identified between virus growth phenotypes on the STE cells and A549 cells and the genetic clades of any genes, either (Figs. S2 and S3).

**Fig. 3.**
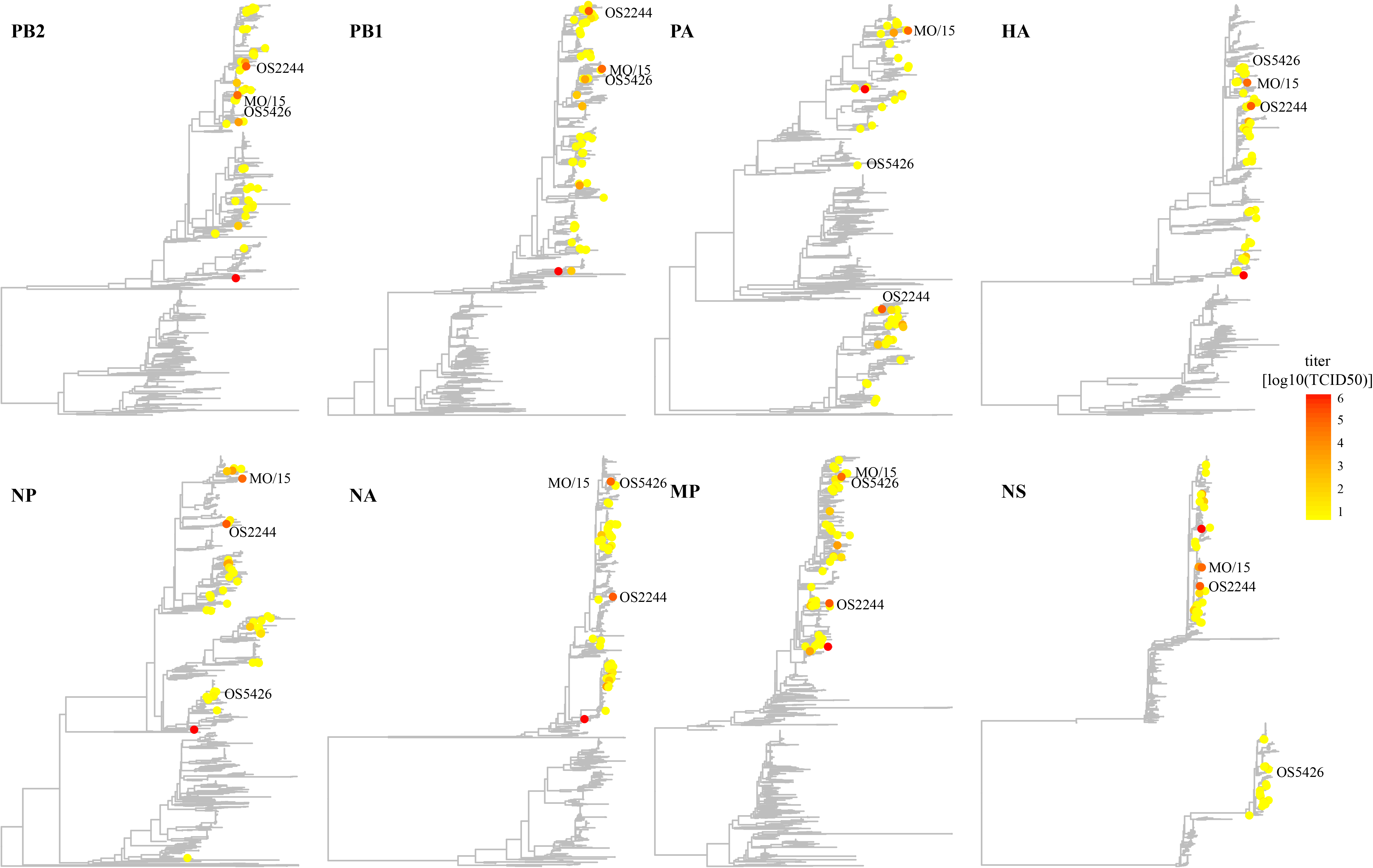
Distribution of growth phenotypic variants in phylogenetic trees. The phenotypes used in this figure was the TCID50 titers on the SNE cells (72 h and 33 °C). The phylogenetic trees for each gene segment was inferred using a maximum-likelihood method implemented in RAxML v8.2.9. A GAMMA model of rate heterogeneity and a generalized time-reversible (GTR) substitution model were applied in the analysis. Phylogenetic trees were then visualized by ggtree v1.6.11. Three viruses, MO/15, OS2244, and OS5425, used in animal study are annotated. Abbreviations are described in the legend of Fig. 1.

### H4N6 avian viruses showed similar receptor binding properties

To evaluate whether the receptor binding properties affect growth phenotypes, we performed molecular analyses of receptor binding sites and receptor binding preference analyses. Sequence analyses suggested that the HA receptor binding sites were conserved across all 115 testing viruses, including OS2244 and OS5426, two prototype viruses we selected in this study. However, compared to avian viruses, both swine isolates, MO/15 and ON/99, had two mutations Q226L and G228S at the HA receptor binding sites (Fig. 4A).

**Fig. 4.**
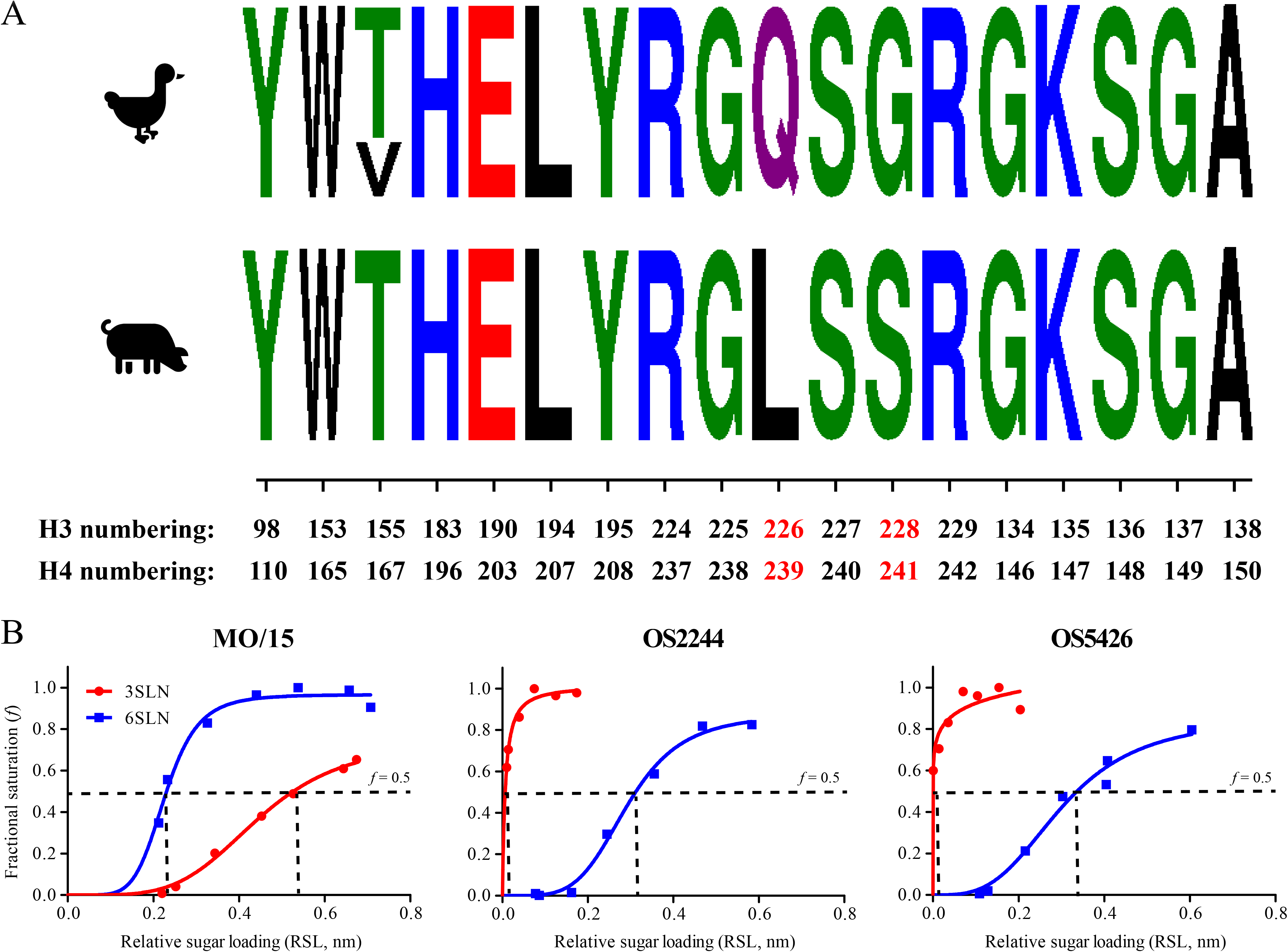
Receptor binding preference of H4N6 IAVs. A) sequence logo of the HA receptor binding sites of avian and swine H4N6 IAVs. Both H3 and H4 numbering were indicated and the mutations at 226 and 228 on HA protein were highlighted. B) receptor binding analyses using biolayer interferometry assays. The 3’-sialyl-N-acetyllactosamine (3′SLN, avian-like IAV receptor analog) and 6’-sialyl-N-acetyllactosamine (6′SLN, human-like IAV receptor analog) were used in these analyses. Streptavidin-coated biosensors were immobilized with biotinylated glycans at different concentrations. Sugar loading– dependent binding signals were captured in the association step and normalized to the same background. Binding curves were fitted by using the binding-saturation method in GraphPad Prism version 7 (https://www.graphpad.com/scientific-software/prism/). Horizontal dashed line indicates half of the fractional saturation (*f* = 0.5), and vertical dashed line indicates relative sugar loading (RSL_0.5_) at *f* = 0.5; the higher the RSL_0.5_, the smaller the binding affinity. Abbreviations are described in the legend of Fig. 1.

The receptor binding specificities were determined through biolayer interferometer assays. The 50% relative sugar loading (RSL_0.5_) were quantified; the smaller the RSL_0.5_, the stronger binding affinity for a testing virus (Fig. 4B). The avian viruses bound to both 3’-sialyl-N-acetyllactosamine (3′SLN, avian-like IAV receptor analog) and 6’-sialyl-N-acetyllactosamine (6′SLN, human-like IAV receptor analog). The avian viruses showed very similar binding patterns with both viruses binding preferentially to 3′SLN rather than to 6′SLN: for 3′SLN, RSL_0.5_ = 0.007 for OS2244 and RSL_0.5_ < 0.001 for OS5426; for 6′SLN, RSL_0.5_ = 0.293 for OS2244 and RSL_0.5_ = 0.303 for OS5426 (Fig. 4B). The swine origin virus (MO/15) also bound to both 3′SLN and 6′SLN but had a higher binding affinity to 6′SLN (RSL_0.5_ = 0.228) than to 3′SLN (RSL_0.5_ = 0.448).

### The ribonucleoprotein (RNP) complex of H4N6 avian viruses showed variations in polymerase activities in mammalian cells

To further identify the molecular mechanisms affecting the diversity in replication phenotypes of H4N6 avian viruses, we determined the polymerase activities of the RNP complex of H4N6 viruses. We hypothesize that the constellation of the RNP complex would affect the polymerase activity of H4N6 IAVs and, thus, the viral replication efficiency. To test this hypothesis, the PB2, PB1, PA, and NP genes of MO/15, OS2244, and OS5246 were cloned and the polymerase activities were quantified using a mini-genome assay in human embryonic kidney (HEK) 293T cells (Fig. 5). Before the mini-genome assay analyses, each gene we cloned was validated to be functional by generating a live reassortant virus with 7 other segments from A/Puerto Rico/8/1934 (H1N1) (abbreviated as PR8) using reverse genetics (data not shown).

**Fig. 5.**
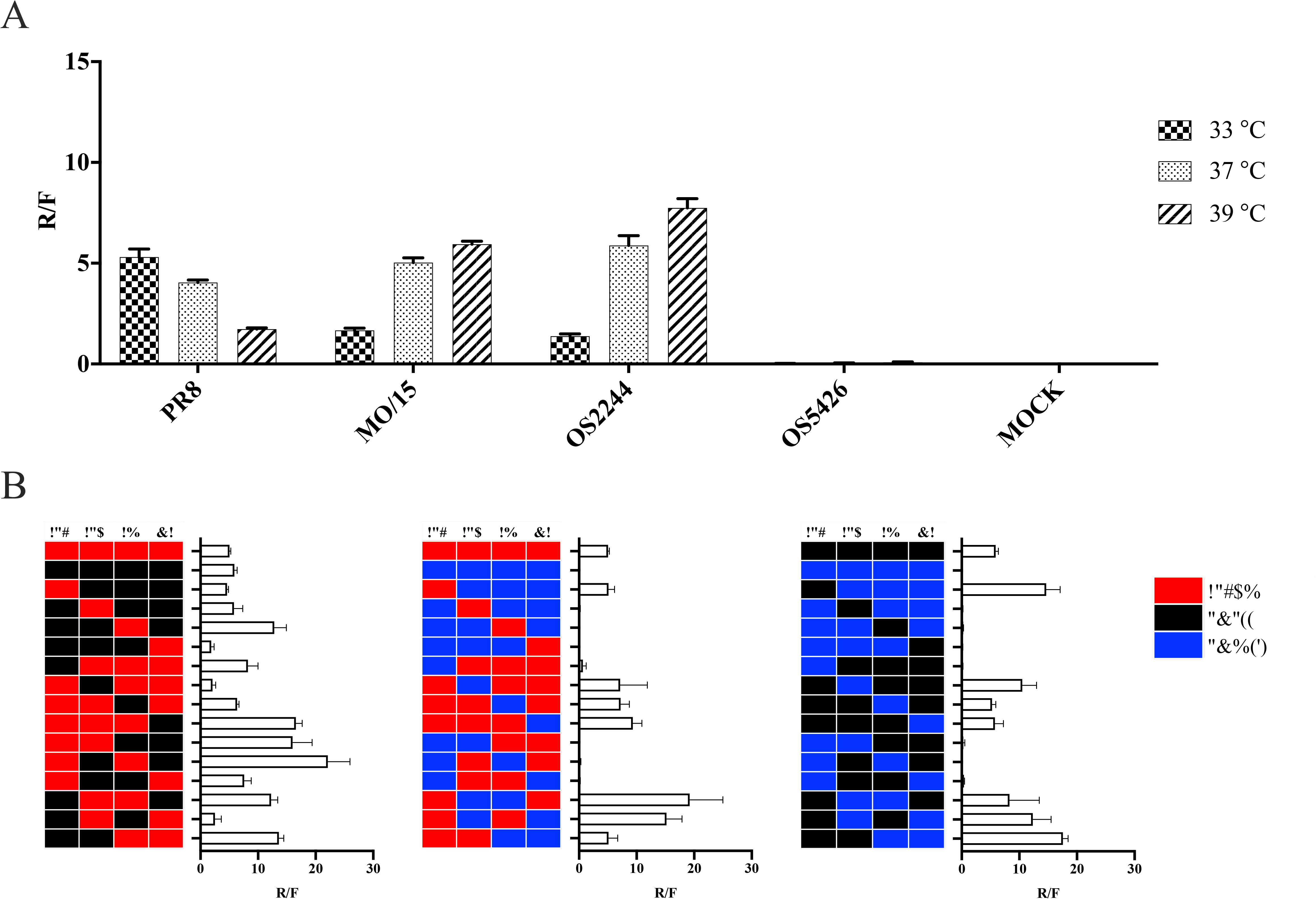
Polymerase activity of the RNP complex. A) The polymerase activities for the RNP complex from wild type viruses on human embryonic kidneys 293T cells at 33, 37 and 39 °C; B) the polymerase activities for the RNP reassortant complex human embryonic kidneys 293T cells at 37 °C. The polymerase activities were determined using minigenome luciferase assays. The mean and standard deviation of the R/F for each RNP complex were derived from the luciferase assay data in triplicate.

Results showed that the polymerase activities for each RNP are consistent with the replication patterns of the corresponding isolates described above. For example, the RNP complex of MO/15 and OS2244 had a R/F value of 1.665 and 1.377 (33 °C), respectively, whereas that of OS5246 was with a R/F < 0.1 (33 °C) (Fig. 5A). In addition, our results showed that the polymerase activities can be temperature dependent (Fig. 5A). For example, MO/15 and OS2244 complexes significantly increased polymerase activity as temperature increased (*p* = 0.0036). As a positive control, polymerase activities of PR8, a laboratory adapted strain, were also quantified. The polymerase activities of PR8 decreased with higher temperature, indicating a difference in the optimal temperature conditions between the RNP complexes from laboratory adapted strains and those from H4N6 wildtype viruses.

In order to investigate the effects of the genomic constellation on the polymerase activities of RNP complex, we determined the polymerase activities of all possible RNP reassortants between MO/15 and OS2244 (n = 16), between MO/15 and OS5426 (n = 16), and between OS2244 and OS5426 (n = 16) (Fig. 5B). Results showed a large extent of diversities among these 48 RNP reassortants (Fig. 5). In general, the polymerase activities of the reassortants between MO/15 and OS2244 are relatively higher than those between OS2244 and OS5426 (*p* < 0.0001) and those between MO/15 and OS5426 (*p* < 0.0001). The polymerase activities between OS2244 and OS5426 and those between MO/15 and OS5426 were not statistically significant (*p* = 0.3315). Further analyses suggested that the PB2 gene is important for high polymerase activity of the RNP complex. All 16 testing RNP complex with OS5246_PB2_ showed a very low or minimal polymerase activity whereas the others with either MO/15_PB2_ or OS2244_PB2_ did not (Fig. 5B).

### The H4N6 swine virus infected pigs and caused limited transmission through direct contacts in pigs

To evaluate the virus infectivity and transmissibility, four pigs (referred to as inoculation pig) were inoculated with MO/15, and three influenza seronegative pigs (contact pigs) were paired with either one or two inoculation pigs in the same pen. Results showed that all four inoculation pigs shed virus in nasal wash samples on 2, 4 and 6 days post-inoculation (dpi) with viral loads ranging from 3.23 (±0.35) to 6.28 (±0.47) Log_10_Copies/mL (Fig. 6 and Table S2), and were further confirmed by reverse-transcription Droplet Digital PCR (RT-ddPCR) with similar viral loads, ranging from 2.79 to 5.89 Log_10_Copies/mL on 2, 4 and 6 dpi (Table S3). To confirm viral infection in the respiratory tracts of pigs, one inoculation pig was euthanized and necropsied on 5 dpi, and another one on 7 dpi. Viral titration by qRT-PCR showed that the upper and middle respiratory tract tissues had viral loads with titers ranging from 4.10 (±0.17) to 5.11 (±0.06) Log_10_Copies/g (Table S4), and the viral loads were further confirmed by RT-ddPCR with titers ranging from 4.29 to 5.21 Log_10_Copies/g (Table S5). Serological analyses by hemagglutination inhibition (HI) assays showed that two remaining inculcation pigs seroconverted as early as on 10 dpi with the same HI titers of 1:160, and remained seroconverted on 21 dpi with HI titers of 1:40 and 1:80, respectively (Table S6).

**Fig. 6.**
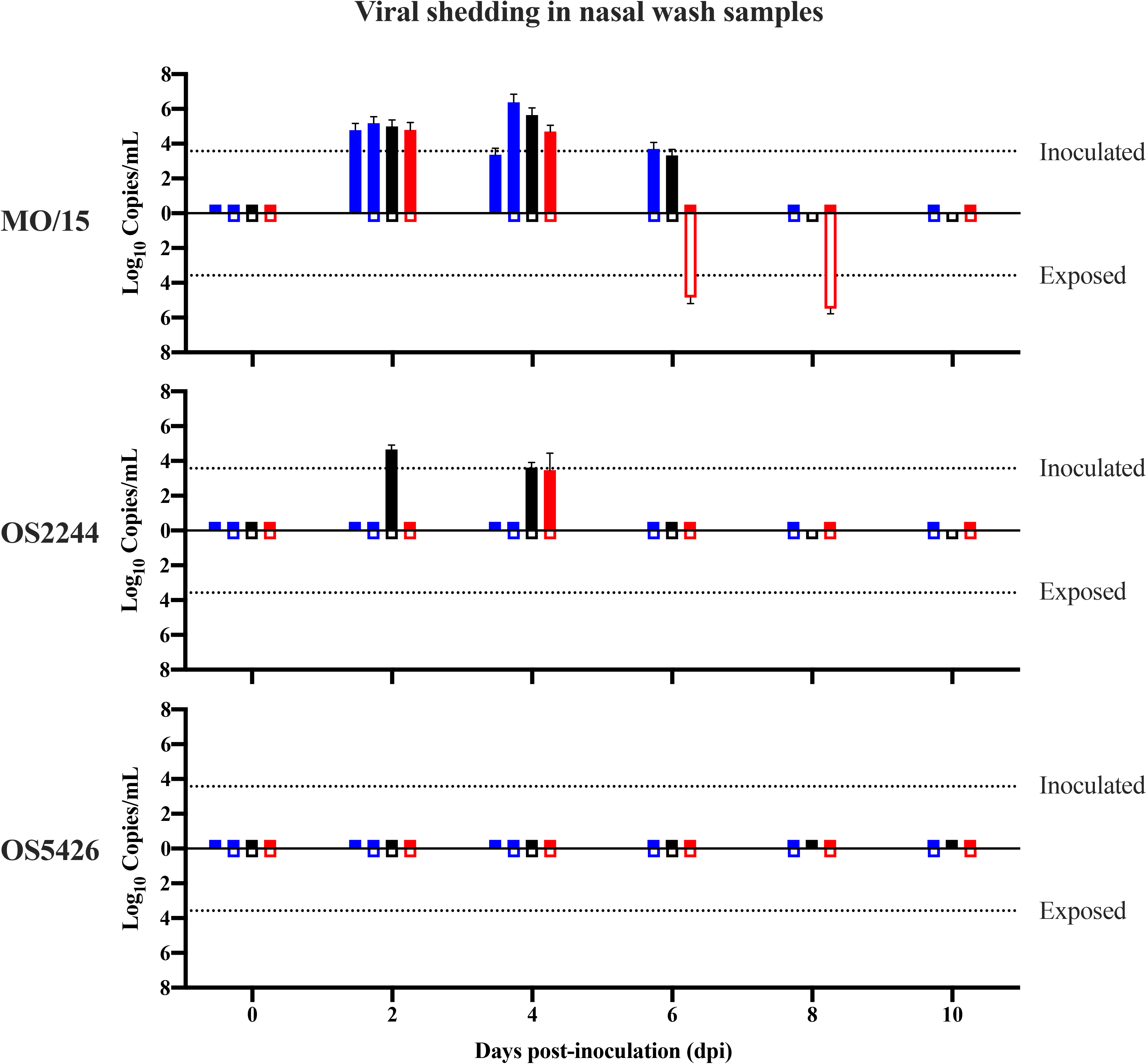
Viral shedding in the animal experiments. For each isolate, seven pigs were assigned into three groups, each with 2 or 3 animals. In each group, one or two animals are used as inoculation animal, and one animal was used as contact animal. Each inoculation animal was inoculated nasally with 10^6^ TCID_50_ of each indicated virus on day 0; the contact animal was brought into the pen 24 h later. Nasal washes were collected at 2-day intervals and one inoculated pig was euthanized and necropsied on day 5 and/or 7, respectively. The same color of bars indicates that the pigs in the same pen. None-filled bars represent inoculation pigs while the filled bars represent contact pigs. The viral titer in each nasal wash sample was determined by qRT-PCR, and the dashed line indicates the limit of detection by qRT-PCR. No viral shedding was detected in all pigs in negative control groups (data not shown).

Among three paired contact pigs, one had virus detected in the nasal wash samples on 6 and 8 dpi with 4.74 (±0.35) and 5.39 (±0.29) Log_10_Copies/mL (Fig 5 and Table S2), respectively. The quantification was also confirmed by RT-ddPCR with similar viral copy numbers, showing 4.40 Log_10_Copies/mL on 6 dpi and 5.06 Log_10_Copies/mL on 8 dpi (Table S3). This shedding contact pig seroconverted on 21 dpi with a HI titer of 1:160 whereas the other two contact pigs remained seronegative against IAV (Table S6).

In summary, our results showed that MO/15 infected feral swine with a high efficiency (4/4; 100%) and that MO/15 caused transmission among pigs through direct contacts, though, with a low transmission efficiency (1/3; 33.33%).

### H4N6 avian IAVs infected pigs but showed distinct phenotypes in virus shedding

We hypothesize that a high replication efficacy in swine respiratory tract epithelial cells, especially the upper respiratory tract epithelial cells, such as SNE, could enhance the viral infectivity and shedding in pigs. To test this hypothesis, we compared the infectivity, virus shedding, and transmission ability in pigs for two prototype avian viruses, OS2244 and OS5426, which showed high and low growth phenotypes respectively in the SNE cells (Fig. 2B). The design of animal experiments is the same as that for MO/15 as described above.

Results showed that two of four pigs inoculated with OS2244 shed viruses in the nasal wash samples with peaking titers of 4.56 (±0.26) Log_10_Copies/mL for pig #156 and 3.38 (±0.97) Log_10_Copies/mL for pig #151 (Fig. 6 and Table S2) and were further confirmed by RT-ddPCR with viral copy numbers ranging from 2.57 to 4.62 Log_10_Copies/mL (Table S3). Viral titrations suggested viruses were detected in the respiratory tract tissues of both the inoculation pigs euthanized on 5 and 7 dpi, with 3.77 (±0.04) to 7.59 (±0.04) Log_10_Copies/g (Table S4) and further confirmed by RT-ddPCR with titers ranging from 3.17 to 8.06 Log_10_Copies/g (Table S5). Both of the two remaining inoculation pigs seroconverted with a HI titer of at least 1:80 (Table S6). These data showed that all four pigs inoculated with OS2244 were infected with the virus and two of these four pigs shed viruses.

In contrast, none of four pigs inoculated with OS5426 had detectable viral loads in all nasal washes collected from 0 through 14 dpi. On the other hand, viral titrations demonstrated viruses were detected in the respiratory tract tissues of the inoculation pig euthanized on 5 dpi (Table S4 and S5). Three remaining pigs inoculated with OS5426 seroconverted with a HI titer ranging from 1:10 to 1:80 (Table S6). These data showed that all four inoculated with OS5426 were infected with the virus but none of them shed viruses.

No viruses were detected in nasal washes from the contact pigs with either OS2244 or OS5466 and those from the negative control pigs; none of the contact pigs with either OS2244 or OS5466 and none of the negative control pigs seroconverted. In addition, none of inoculation pigs, contact pigs, and negative control pigs showed any clinical signs.

In summary, the animal study showed that OS2244, which replicated efficiently in SNE, shed viruses in their nasal wash samples whereas OS5426, which replicated poorly in SNE, did not shed any viruses in their nasal wash samples, validating our hypotheses, although both avian viruses effectively infected pigs through nasal inoculation.

## Discussion

In this study, through phenotypic analyses of 115 genetically diverse H4N6 avian IAVs, we identified only a small set of isolates with a high growth efficiency on the epithelial cells of the upper swine respiratory tract (i.e. SNE). A similar high growth phenotype was also identified in a H4N6 isolate that was detected in a spillover event from wild birds to domestic swine. Genetic analyses suggested that the genetic constellation of the RNP complex, but not the receptor binding properties, be a major factor contributing to the observed phenotypic diversity. Animal studies suggested that both of the viruses with a high growth phenotype on SNE cells as well as the virus with a low growth phenotype on SNE cells infected pigs through nasal inoculation. Only the pigs infected with the high growth phenotypes had detectable viral shedding. As shown in a simplified model, those viruses with high growth phenotypes on SNE cells could potentially enable virus to be transmitted to other pigs, thus helping the virus acquire adaptive mutations or a gene from other co-circulating viruses through genetic reassortment (Fig. 7). For example, Q226L and G228S were detected in HA protein of both H4N6 swine isolates but rarely in avian isolates, and these mutations increased human-like receptor binding ability and could have further enhanced virus transmission ability in pigs (Fig. 4) [33, 34]. In summary, this study suggested that tissue tropisms of H4N6 avian IAVs affect their spillovers from wild birds to pigs. Thus, characterization of the tissue tropism of avian IAVs could be an efficient way for risk assessment, such as in evaluating their potentials to be transmitted from avian to pigs.

**Fig. 7.**
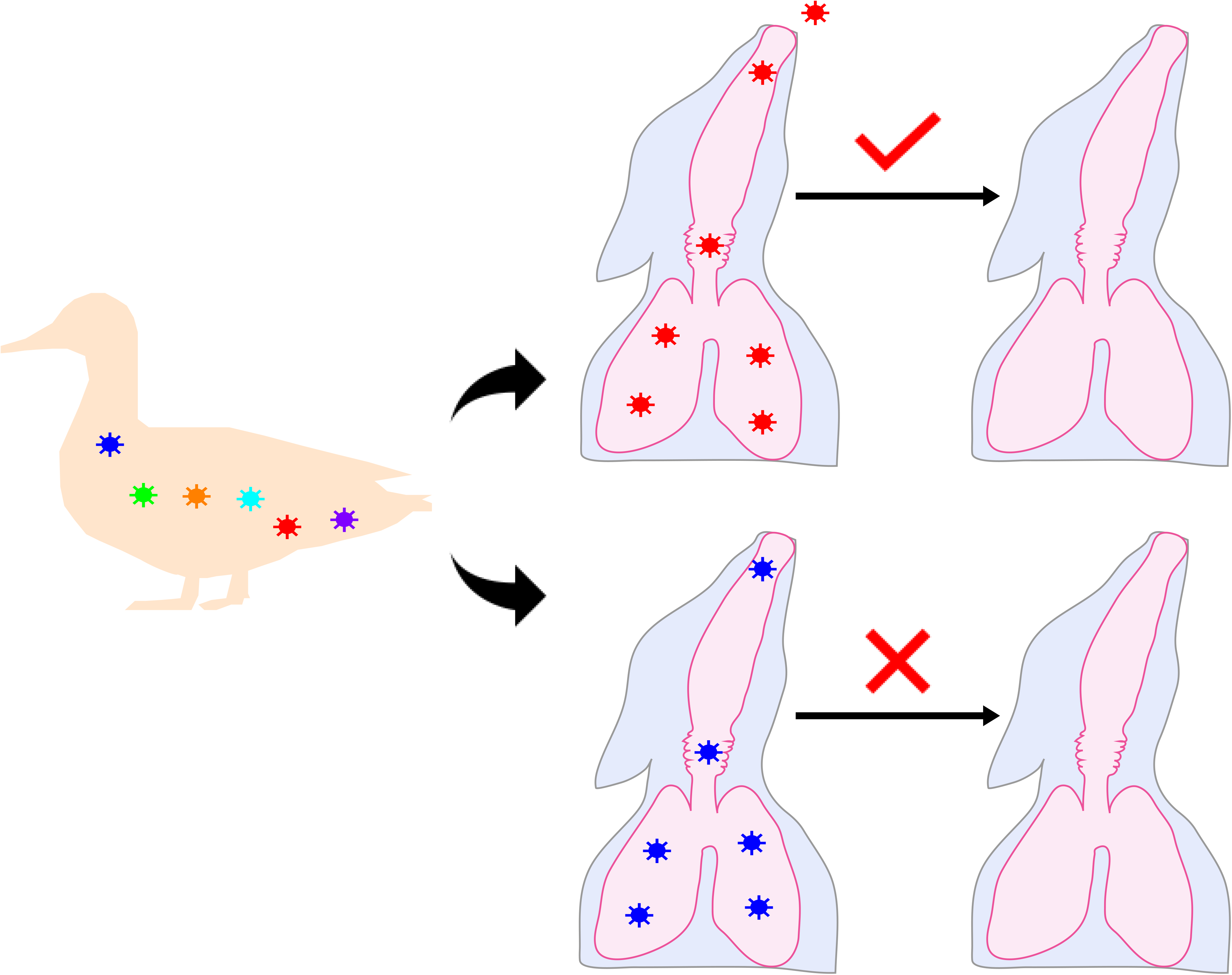
A simplified transmission model for avian IAVs from wild birds to pigs. The wild aquatic birds maintain a large pool of genetically and phenotypic diverse IAVs, and only a few of these viruses can replicate well in the swine upper respiratory tract epithelial cells. Although many of these viruses can infect pigs, only the viruses which can replicate well in the swine upper respiratory tract epithelial cells can cause virus shedding in pigs upon infection, providing opportunities for these viruses to become adapted to pig, thus posing a higher potential risk for creating novel variants or detrimental reassortants within pig populations.

In wild birds, there is a lack of selection pressure for mammal-based replication efficiency. While wild birds maintain a large genetically diverse pool of IAVs, the majority of wild type viruses would not be expected to bear a high growth phenotype in mammalian cells. Our results from screening 115 H4N6 avian viruses showed that only four reached the same growth titers of MO/15 in SNE cells and that the high growth viruses are scattered across the phylogenetic trees of eight genetic segments (Fig. 2). No particular genetic constellations correlated with the varied growth phenotypes we determined (Figs. 3, S2, and S3). Our study suggests that risk assessment of avian IAVs shall include phenotypic analyses of a broad range of genetically diverse isolates rather than only the isolates in those most commonly found genotypes. In prior studies, a few genetic markers, such as 627 in PB2 [35–37], and 701 in PB1 [36], were shown to increase virus replication efficiencies in mammals, such as mice and ferrets. Thus, it would be useful to develop a genomic sequence-based risk assessment tool by identifying synergistic genetic markers through a large set of phenotypic data.

Among all subtypes of avian IAVs, subtype H4 IAVs are enzootic in both wild birds and domestic poultry and may be exposed to both pigs and humans [3, 24, 25, 38–42]. In addition to two spillovers of H4N6 IAVs mentioned above in Ontario, Canada (1999) [26] and in Missouri, USA (2015) [24], H4N1 and H4N8 isolates were recovered from pigs in China [43, 44]. Because of long term co-circulation with multiple subtypes of IAVs in wild aquatic birds, H4 avian IAVs had frequent reassortments and showed a large extent of genotypic diversity [39]. Prior studies have also shown that these H4 avian viruses maintain a large pool of phenotypic diversity. For example, some H4 isolates exhibited infectivity in pigs and mice without prior adaptation and caused efficient transmission in guinea pigs [11, 39]. Similar to the receptor binding properties of two H4N6 avian viruses we demonstrated in this study (Fig. 4), these avian H4 IAVs bind to both avian-like and human-like receptors with a stronger affinity to avian-like receptors [39]. On the other hand, similar to our results (Fig. 5), another study suggested that the RNP complex of H4 IAVs contributed to higher virulence in mice [45]. These data support our findings that, in addition to glycan receptor binding, the genomic makeup, such as those in the RNP complex, can affect virus infectivity and transmissibility of avian IAVs in mammals.

The animal study demonstrated that the swine H4N6 virus, MO/15, shed viruses and also caused limited transmission (33%) in pigs through direct contacts (Fig. 6). This result is contradictory to the results reported in a prior study that showed that the same isolate did not cause virus transmission in pigs [24]. To validate our analyses, we performed viral titration using TCID50 in viral sheds among those four inoculation pigs and one contact pigs in the animal experiment with MO/15 (Table S2 and S3). Our results confirmed that 3 of 4 pigs inoculated with MO/15 had viral titers ranging from 2.50 to 3.67 Log_10_TCID_50_/mL and the contact pig with a viral titer of 4.5 Log_10_TCID_50_/mL at 8 dpi (data not shown). The nasal washes in one of four inoculation pigs did not have detectable TCID50, possibly due to a relatively lower number of viral loads and the limited growth preference of avian viruses on MDCK cells. Compared with domestic pigs used in the other study, the feral swine used in this study are less likely to be primed with IAVs [46] although the pigs in both studies were shown to be seronegative against IAVs. On the other hand, the inoculation routes and doses are different in these two studies. In this study, 10^6^ TCID_50_ in 2 mL of inoculum (1 mL per nostril) was intranasally inoculated for each pig; in the other study, a relatively lower concentration (3 × 10^5^ TCID_50_ in 3 mL) of virus was inoculated into each pig intranasally (1 mL of virus) and intratracheally (2 mL of virus) [24]. It has been shown that viral loads, inoculation volume, and inoculation routes can affect virus infection ability [47]. Nevertheless, the H4N6 swine isolate still seems to have limited transmission ability in pigs, which is consistent with the low seroprevalence among the pigs at the outbreak farm and the other pigs in the production system [24].

In summary, this study suggested that a large pool of phenotypic variants exist among avian IAVs, only a few of which may cause virus shedding in swine upon infection, providing more opportunities for these avian viruses to become pig adapted, thus posing a higher potential risk for creating novel variants or detrimental reassortants within swine populations.

## Materials and Methods

### Cells

Madin-Darby canine kidney (MDCK) cells, Human embryonic kidney (HEK) 293T cells and human alveolar adenocarcinoma (A549) cells (American Type Culture Collection, Manassas, VA, USA) were maintained in Dulbecco’s modified Eagle’s medium (DMEM, Gibco, New York, USA) supplemented with 10% fetal bovine serum (FBS; Atlanta Biologicals, Lawrenceville, GA, USA) at 37°C under 5% CO_2_. Swine nasal epithelial cells (SNE) and swine trachea epithelial cells (STE) were kindly provided by Dr. Stacey Schultz-Cherry, St Jude Children’s Research Hospital and grown at 37 °C with 5% CO_2_ in DMEM/F12 (Thermo Fisher Scientific, Asheville, CA) supplemented with FBS (10%).

### Viruses

A/swine/Missouri/A01727926/2015 (H4N6) (abbreviated as MO/15), an isolate recovered from a sick domestic swine, was kindly provided by the USDA Swine Influenza Surveillance Program and propagated for one passage on MDCK cells at 37°C with 5% CO_2_ in Opti-MEM I Reduced Serum Medium (Thermo Fisher Scientific, Asheville, NC, USA) supplemented with 1 μg/mL of TPCK-trypsin (Gibco, New York, USA) before being used in the molecular studies, phenotypic determination and the animal study.

A total of 115 H4N6 wild bird origin isolates from North America were selected to represent genomic diversity in addition to the diversity in avian hosts, sampling location, and sampling time (Table S1). These avian isolates were propagated for one passage in specific pathogen-free (SPF) 10-day-old chicken embryonated eggs (Charles River Laboratories, Inc., Norwich, CT) before being used in this study for the molecular studies, phenotypic determination, and the animal study.

### Hemagglutination and hemagglutination inhibition assays

Hemagglutination and hemagglutination inhibition (HI) assays were carried out by using 0.5% turkey erythrocytes as previously described [48].

### Viral titration

For viral titration, the 50% tissue culture infection dose (TCID_50_) was determined on MDCK cells. Briefly, MDCK cells were seeded in 96-well plate at 2 × 10^4^/well with Opti-MEM I Reduced Serum Medium. Cells were incubated at 37°C with 5% CO_2_ for 18-20 h before virus inoculation. Viral samples were serial diluted in Opti-MEM I Reduced Serum Medium supplemented with 1 μg/mL of TPCK-trypsin. Cell medium was removed and 100 μL of each virus dilution was inoculated onto MDCK cells in quadruplicate. Infected cells were incubated at 37°C with 5% CO_2_ for 72 h and then the number of positive and negative wells for each dilution were recorded for TCID_50_ calculation. The TCID_50_ was calculated as previously described by Reed and Muench [49].

### Molecular cloning and genomic sequencing

The viral RNA was extracted using GeneJet Viral DNA/RNA extraction kit (Thermo Fisher Scientific, Asheville, NC, USA) and cDNA was transcript upon the viral RNA as template using SuperScript™ III Reverse Transcriptase (Invitrogen, Thermo Fisher Scientific, Asheville, NC, USA). The PB2, PB1, PA, and NP gene of MO/15, OS2244, and OS5426 were cloned into pHW2000 vector with universal primers [50]. After cloning, the plasmids were prepared for sequencing to further confirm the sequences were correct and no additional mutations.

### Minigenome

For luciferase assay, 293T cells were plated in 96-well plate at 4 × 10^4^/well for transfection on the next day when cell density reached 80% of confluence. The 293T cells were transfected with 40 ng of each RNP gene plasmids pHW2000-PB2, pHW2000-PB1, pHW2000-PA, and pHW2000-NP, 40 ng of Human r-Luc-*Renilla* Luciferase reporter plasmid, and 4 ng of pGL4.13-Firefly Luciferase reporter plasmid using Lipofectamine 2000 (Invitrogen, Carlsbad, CA) in Opti-MEM (Gibco, Carlsbad, CA) for 48 h. Transfection was performed in triplicate. The luciferase activity was developed using the dual-luciferase reporter system (Promega, Madison, WI) according to the manufacturer’s instructions and measured on a Cytation 5 (BioTek, Winooski, VT).

### Replication efficiency

To determine the replication efficiency of H4N6 viruses, we determined the viral titers for these viruses after they are amplified for 72 h in SNE cells at 33 °C, STE cells at 37 °C, and A549 cells at 39 °C.

### Growth kinetics

The growth kinetics were determined in SNE, STE, and A549. Briefly, cells were seeded in 6-well plates at a density of 5 × 10^5^ cells/well. Twenty-four hours later, cells were washed once with PBS and then infected with a testing virus at a multiplicity of infection (MOI) of 10^-3^ TCID_50_/cell. After absorption for 1 h at 37 °C, the inoculum was removed. The cells were washed once with PBS and then 3 mL of Opti-MEM supplemented with 1 μg/mL of TPCK-trypsin was added. Cultures were incubated at 33 °C, 37 °C, or 39 °C for the duration of the experiment. At 12, 24, 48, and 72 h post-infection, the supernatants were collected, and their TCID_50_ were then determined. In addition to MO/15, nine representative avian isolates (Table S1) with distinct growth phenotypes across three testing temperatures were used for the growth kinetics analyses.

### RNA extraction and viral copy number determination

Viral RNA was extracted from nasal wash and tissue samples by using the MagMAX Pathogen RNA/DNA Kit (Thermo Fisher Scientific, Asheville, NC, USA) according to the manufacturer’s instructions. Quantitative reverse-transcription PCR (qRT-PCR) was used to determine the copy numbers of IAVs in nasal wash samples and tissue homogenized samples as previously described [51]. Briefly, qRT-PCR was performed in triplicate by using TaqMan Fast Virus 1-step Master Mix (Life Technology, Carlsbad, CA) with the M gene specific primer set: Forward 5’-GACCRATCCTGTCACCTCTGAC -3’; Reverse 5’-AGGGCATTYTGGACAAAKCGTCTA -3’; and FAM-Probe 5’-TGCAGTCCTCGCTCACTGGGCACG -3’. Viral copies in samples were determined with the standard curve, which was generated by the plasmid containing the M gene of PR8. The copy number determination by qRT-PCR were performed in triplicate.

Quantification of viral copies in both nasal wash and tissue homogenize samples were confirmed by using reverse-transcription droplet digital PCR (RT-ddPCR) on a QX200TM Droplet Digital PCR system (Bio-Rad, Hercules, CA) with One-Step RT-ddPCR Advanced Kit for Probes (Bio-Rad, Hercules, CA). The influenza A-specific primers and probe set used in RT-ddPCR are the same as those used in qRT-PCR. The experiment of RT-ddPCR was performed in duplicate and the data was shown as the average of the two values.

### Viral purification and quantification of virus particles

Viruses were purified by sucrose gradient centrifugation as describe elsewhere [23]. The purified viruses were dissolved in PBS and dialyzed against PBS at 4°C overnight. The concentrations of virus particles were determined using sodium dodecyl sulfate–polyacrylamide gel electrophoresis as described elsewhere [52].

### Virus-glycan receptor binding assay and data analyses

Two biotinlyated glycan analogs, carbohydrates 3’-sialyl-N-acetyllactosamine (3′SLN) representing SA2,3GA and 6’-sialyl-N-acetyllactosamine (6′SLN) representing SA2,6GA, were purchased from GlycoTech (Gaithersburg, MD). The glycan stocks were reconstituted at 1 mg/ml in 50% glycerol - PBS (vol/vol) solution according to the manufacturer’s instructions and were stored at 4°C until use. Binding of viruses (at the concentration of 5 nM) to the biotinylated glycan analogs was performed as previously described in an Octet RED96 biolayer interferometer equipped with streptavidin biosensor tips (PALL FortéBIO, Menlo Park, CA, USA). The glycan concentrations ranged from 0.007 to 1.5 μg/mL.

Responses were normalized by the highest value obtained during the experiment, and binding curves were fitted by using the binding-saturation method in GraphPad Prism version 8 (https://www.graphpad.com/scientific-software/prism/). The normalized response curves report the fractional saturation (*f*) of the sensor surface as described in a previous study [53]. RSL_0.5_ (relative sugar loading, *f* = 0.5) was used to quantitate the binding affinity of two selected viruses against two glycan analogs. The higher the RSL_0.5_, the weaker the binding affinity.

### Feral swine

For animal experiments, a total of 26 feral swine (body weight 16–22 kg) were trapped in a rural area of Starkville, MS, USA, by using corral traps similar to those previously described [54]. Animals were transported to the National Wildlife Research Center, Mississippi Field Station, in Mississippi State, MS, USA, where they were quarantined for 1 week. Before the animals were included in the experiments, we confirmed that these feral swine had not been exposed to brucellosis, pseudorabies and IAV using ELISA ((IDEXX, Westbrook, ME) as previously described [54] and that all HI assay results were negative for three testing H4N6 IAVs (MO/15, OS2244, and OS5426), and three endemic human IAVs [i.e., A/California/04/2009(H1N1), A/Switzerland/9715293/2013(H3N2), and A/Hong Kong/4801/2014(H3N2)]. The swine were then housed and fed according to standard protocol, as described elsewhere [54].

### Animal experiment for evaluating the pathogenesis and transmission of H4N6 IAVs

Twenty-six feral swine were randomly assigned into four experimental groups, including 3 treatment groups and 1 control group. Treatment groups included 7 pigs each and were exposed to one of three testing H4N6 isolates, MO/15, OS2244, and OS5426.

Pigs in each treatment group were housed in 3 pens (2 or 3 pigs/pen), and pigs in the control groups in 2 pens (2 or 3 pigs/pen). Within each pen, 1 or 2 pigs were experimentally inoculated with virus or sterile PBS while the remaining co-housed pig served as the contact pig. The control group was housed in a separate building from the experimental groups. Prior to virus inoculation and sample collection, pigs were anesthetized using a method described previously [54]. After being anesthetized, four pigs within each treatment group were intranasally inoculated with 10^6^ TCID_50_ of virus in a volume of 2 mL (1 mL/nostril). Three IAV seronegative contact pigs were co-housed 24 hours post virus inoculation. Two pigs in the control group were intranasally inoculated with 2 mL of PBS.

On 1-10 dpi, nasal wash fluids were collected from both nostrils of all pigs into 3 mL of PBS with 100 unit/mL of Pen-Strep and then subjected to viral copy number determination and titration by TCID_50_. The body temperature of each pig was measured before samples were taken. Serum from each pig was also collected for seroconversion analysis using an HI assay with 0.5% turkey red blood cells, as described previously [48]. At 5 and 7 dpi, respectively, one pig from each treatment group and one control pig were euthanized and necropsy was performed according to a previously described procedure [54].

Tissues of respiratory tract were collected for viral titration: left cranial lung (LCR), left caudal lung (LCD), right cranial lung (RCR), right caudal lung (RCD), right middle lung (RMD), right accessory (RA), upper trachea (TR-U), middle trachea (TR-M), distal trachea (TR-D), bronchus (BR), soft palate (SP), ethmoid turbinate (ET), rostral turbinate (RT), and middle turbinate (MT). To quantify IAV, each tissue was homologized with the ratio of 1:4 (weight/volume; 1 gram tissue in 4 mL of buffer) in PBS containing 100 unit/mL of Pen-Strep and then subjected the solutions to three freeze-thaw cycles prior to performing RNA extraction and virus titrations.

### Genomic sequences

A total of 1,497 H4N6 genomic sequences were downloaded from Influenza Research Database (IRD), Global Initiative on Sharing All Influenza Data (GISAID), and National Center for Biotechnology Information (NCBI) on May 2, 2017, and the genomic sequences of 35 H4N6 isolates were obtained from USDA. The repeated sequence records were removed and data from the three databases were merged together. A total of 115 avian H4N6 isolates from North America were selected to represent the wide genomic diversity of H4N6 IAVs in North America as well as to a diversity of avian host species, sampling times, and sampling locations (Table S1).

### Multiple sequence alignment and phylogenetic analysis

Multiple sequence alignments (MSA) were generated by using MAFFT v7.273 [55]. The phylogenetic analysis was performed using a maximum-likelihood tree to represent the evolutionary relationship among different isolates, as described elsewhere [56, 57]. The phylogenetic trees for all gene segments (PB2, PB1, PA, HA, NP, NA, MP, and NS) were inferred using a maximum-likelihood method implemented in RAxML v8.2.9 [58]. A GAMMA model of rate heterogeneity and a generalized time–reversible (GTR) substitution model were applied in the analysis. Phylogenetic trees were then visualized by ggtree v1.6.11 [59].

### Statistical analyses

The nonparametric method, Kruskal-Wallis test (GraphPad Prism, https://www.graphpad.com/scientific-software/prism/), was used to test the hypothesis that the polymerase activities of influenza viruses would be affected by temperature. This analysis was performed on the polymerase activities of RNP_MO/15_, RNP_OS2244_, and RNP_PR8_ under three testing temperatures, 33, 37, and 39 °C. To test the hypothesis that the genomic constellation would affect the polymerase activities of the RNP complex, a 2-way ANOVA was used to compare the polymerase activity difference between RNP^MO/15×OS2244^ and RNP^MO/15×OS5426^, or RNP^MO/15×OS2244^ and RNP^OS2244×OS5426^, or RNP^MO/15×OS5426^ and RNP^OS2244×OS5426^. A p-value of 0.05 was determined statistically significant for all analyses.

### Biosafety protocol for laboratory and animal experiments

Virus titration and purification and virus inoculation in feral swine were conducted under Biosafety Level 2 conditions, in compliance with U.S. Department of Agriculture–approved protocols of Institutional Biosafety Committee.

### Ethics statement

The animal experiments were performed under the protocol numbers QA2625 titled Potential of Avian influenza A virus to infect feral swine, which was approved by the Institutional Animal Care and Use Committee of National Wildlife Research Center (NWRC) in accordance with the USDA Animal Welfare Regulations.

## Acknowledgments

We would like to acknowledge the USDA Swine Influenza Surveillance Program for providing influenza virus isolate and Dr. Stacy at St. Jude for providing swine primary cells. This project was partially supported by grant R21AI135820 from the National Institutes of Health.

## Supplementary Information

### Supplementary Tables

**Supplementary Table 1.** List of H4N6 viruses used in this study. The passages in specific pathogen free embryonated chicken eggs or MDCK cells are annotated, and the HA titers were determined using 0.5 turkey red blood cells. The TCID50 was determined in MDCK cells.

| Virus Name | Subtype | Passage | HA Titer | Log<sub>10</sub>TCID<sub>50</sub>/mL |
| --- | --- | --- | --- | --- |
| A/Swine/Missouri/A01727926/2015 | H4N6 | MDCK3 | 2<sup>7</sup> | 6.67 |
| A/blue-winged teal/Ohio/15OS5426/2015 | H4N6 | EGG1 | 2<sup>9</sup> | 5.5 |
| A/mallard/Ohio/08OS1270/2008 | H4N6 | EGG1 | 2<sup>9</sup> | 7.33 |
| A/blue_winged_teal/Illinois/10OS1561/2010 | H4N6 | EGG1 | 2<sup>9</sup> | 7 |
| A/American_green_winged_teal/Mississippi/11OS5903/2011 | H4N6 | EGG1 | 2<sup>8</sup> | 6.33 |
| A/blue_winged_teal/Ohio/12OS2106/2012 | H4N6 | EGG1 | 2<sup>8</sup> | 7.33 |
| A/mallard/Wisconsin/11OS4309/2011 | H4N6 | EGG1 | 2<sup>10</sup> | 6.67 |
| A/blue_winged_teal/Ohio/12OS2244/2012 | H4N6 | EGG1 | 2<sup>9</sup> | 7.5 |
| A/blue_winged_teal/Ohio/12OS2293/2012 | H4N6 | EGG1 | 2<sup>9</sup> | 6.5 |
| A/blue_winged_teal/Missouri/11OS2620/2011 | H4N6 | EGG1 | 2<sup>8</sup> | 6.67 |
| A/environment/Maryland/1655/2006 | H4N6 | EGG1 | 2<sup>8</sup> | 7.67 |
| A/environment/Maryland/09OS1315/2009 | H4N6 | EGG1 | 2<sup>8</sup> | 7.67 |
| A/mallard/Maryland/10OS1040/2010 | H4N6 | EGG1 | 2<sup>9</sup> | 8.5 |
| A/mallard/Wisconsin/1538/2009 | H4N6 | EGG1 | 2<sup>9</sup> | 7.5 |
| A/Americangreen-wingedteal/Ohio/13OS1429/2013 | H4N6 | EGG1 | 2<sup>9</sup> | 7.5 |
| A/blue-wingedteal/Ohio/12OS2295/2012 | H4N6 | EGG1 | 2<sup>9</sup> | 7.67 |
| A/mallard/Maryland/09OS1134/2009 | H4N6 | EGG1 | 2<sup>9</sup> | 7.67 |
| A/mallard/Maryland/11OS951/2011 | H4N6 | EGG1 | 2<sup>9</sup> | 8.33 |
| A/blue-wingedteal/Missouri/10MO003/2010 | H4N6 | EGG1 | 2<sup>9</sup> | 8 |
| A/blue-wingedteal/Iowa/13OS2359/2013 | H4N6 | EGG1 | 2<sup>9</sup> | 7 |
| A/mallard/Wisconsin/11OS3481/2011 | H4N6 | EGG1 | 2<sup>8</sup> | 8 |
| A/mallard/Maryland/08OS1421/2008 | H4N6 | EGG1 | 2<sup>8</sup> | 7.5 |
| A/mallard/Wisconsin/2653/2009 | H4N6 | EGG1 | 2<sup>8</sup> | 8 |
| A/Americangreen-wingedteal/Ohio/13OS2050/2013 | H4N6 | EGG1 | 2<sup>8</sup> | 7.67 |
| A/environment/Maryland/09OS1301/2009 | H4N6 | EGG1 | 2<sup>8</sup> | 7.67 |
| A/northernshoveler/Illinois/11OS4707/2011 | H4N6 | EGG1 | 2<sup>9</sup> | 8 |
| A/mallard/Maryland/11OS964/2011 | H4N6 | EGG1 | 2<sup>9</sup> | 8.5 |
| A/mallard/Maryland/14OS1477/2014 | H4N6 | EGG1 | 2<sup>8</sup> | 7.67 |
| A/mallard/Maryland/09OS1170/2009 | H4N6 | EGG1 | 2<sup>6</sup> | 6.67 |
| A/environment/Maryland/09OS1302/2009 | H4N6 | EGG1 | 2<sup>8</sup> | 7.67 |
| A/mallard/Maryland/14OS1322/2014 | H4N6 | EGG1 | 2<sup>7</sup> | 6.33 |
| A/mallard/Iowa/10OS2420/2010 | H4N6 | EGG1 | 2<sup>8</sup> | 7.67 |
| A/blue-wingedteal/Ohio/12OS2321/2012 | H4N6 | EGG1 | 2^8 | 7 |
| A/blue-wingedteal/Ohio/12OS3128/2012 | H4N6 | EGG1 | 2^8 | 7.5 |
| A/green-wingedteal/Ohio/1292/2005 | H4N6 | EGG1 | 2^9 | 8 |
| A/mallard/Maryland/08OS1414/2008 | H4N6 | EGG1 | 2^8 | 7.67 |
| A/mallard/Ohio/1695/2009 | H4N6 | EGG1 | 2^8 | 7.67 |
| A/northernpintail/Wisconsin/11OS3299/2011 | H4N6 | EGG1 | 2^7 | 7.5 |
| A/northernpintail/Ohio/15OS5731/2015 | H4N6 | EGG1 | 2^8 | 8 |
| A/environment/Maryland/14OS1379/2014 | H4N6 | EGG1 | 2^9 | 6.67 |
| A/Americangreen-wingedteal/Ohio/13OS2063/2013 | H4N6 | EGG1 | 2^8 | 8.5 |
| A/blue-wingedteal/Wisconsin/11OS3027/2011 | H4N6 | EGG1 | 2^8 | 8.33 |
| A/redhead/Ohio/13OS0376/2013 | H4N6 | EGG1 | 2^8 | 8.33 |
| A/blue-wingedteal/Ohio/15OS5173/2015 | H4N6 | EGG1 | 2^6 | 6.67 |
| A/mallard/Wisconsin/10OS2918/2010 | H4N6 | EGG1 | 2^8 | 7.67 |
| A/mallard/Ohio/660/2002 | H4N6 | EGG1 | 2^6 | 6 |
| A/mallard/Maryland/07OS1669/2007 | H4N6 | EGG1 | 2^8 | 7.5 |
| A/mallard/Ohio/657/2002 | H4N6 | EGG1 | 2^8 | 8 |
| A/mallard/Ohio/15OS4260/2015 | H4N6 | EGG1 | 2^9 | 7.67 |
| A/blue-wingedteal/Ohio/12OS4593/2012 | H4N6 | EGG1 | 2^8 | 7.5 |
| A/mallard/Ohio/14OS3201/2014 | H4N6 | EGG1 | 2^8 | 7.5 |
| A/environment/Maryland/09OS1293/2009 | H4N6 | EGG1 | 2^8 | 6.67 |
| A/environment/Maryland/09OS1314/2009 | H4N6 | EGG1 | 2^8 | 7.67 |
| A/environment/Maryland/2115/2006 | H4N6 | EGG1 | 2^5 | 6.33 |
| A/blue-wingedteal/Ohio/1850/2006 | H4N6 | EGG1 | 2^9 | 7.67 |
| A/Americangreen-wingedteal/Ohio/13OS2021/2013 | H4N6 | EGG1 | 2^9 | 8 |
| A/mallard/Ohio/671/2002 | H4N6 | EGG1 | 2^8 | 7.33 |
| A/mallard/Ohio/10OS1324/2010 | H4N6 | EGG1 | 2^8 | 7 |
| A/environment/Maryland/09OS1291/2009 | H4N6 | EGG1 | 2^8 | 7.5 |
| A/green-wingedteal/Ohio/1324/2005 | H4N6 | EGG1 | 2^8 | 8 |
| A/mallardduck/Ohio/15OS5358/2015 | H4N6 | EGG1 | 2^7 | 5.5 |
| A/mallard/Maryland/14OS1312/2014 | H4N6 | EGG1 | 2^8 | 6.33 |
| A/mallard/Maryland/14OS1310/2014 | H4N6 | EGG1 | 2^8 | 6.5 |
| A/mallard/Illinois/08OS2315/2008 | H4N6 | EGG1 | 2^8 | 8 |
| A/blue-wingedteal/Missouri/15OS4858/2015 | H4N6 | EGG1 | 2^8 | 7.5 |
| A/mallard/Ohio/11OS2085/2011 | H4N6 | EGG1 | 2^8 | 7.67 |
| A/mallard/Ohio/15OS3997/2015 | H4N6 | EGG1 | 2^8 | 8.33 |
| A/Americangreen-wingedteal/Illinois/10OS1551/2010 | H4N6 | EGG1 | 2^9 | 7.5 |
| A/environment/Maryland/09OS1292/2009 | H4N6 | EGG1 | 2^8 | 7.67 |
| A/mallard/Ohio/298/1987 | H4N6 | EGG1 | 2^9 | 7.5 |
| A/mallard/Ohio/1501/2006 | H4N6 | EGG1 | 2^6 | 7.33 |
| A/mallard/Ohio/15OS4235/2015 | H4N6 | EGG1 | 2^9 | 7.5 |
| A/blue-wingedteal/Missouri/10MO0407/2010 | H4N6 | EGG1 | 2^7 | 7.5 |
| A/blue-wingedteal/Wisconsin/2753/2009 | H4N6 | EGG1 | 2^8 | 7 |
| A/Americangreen-wingedteal/Ohio/12OS2137/2012 | H4N6 | EGG1 | 2^8 | 7.33 |
| A/mallard/Ohio/15OS4181/2015 | H4N6 | EGG1 | 2^7 | 4.5 |
| A/mallard/Ohio/15OS3981/2015 | H4N6 | EGG1 | 2^8 | 4.5 |
| A/blue-wingedteal/Missouri/10MO030/2010 | H4N6 | EGG1 | 2^8 | 8 |
| A/environment/Maryland/09OS1290/2009 | H4N6 | EGG1 | 2^8 | 7.67 |
| A/blue-wingedteal/Missouri/15OS4861/2015 | H4N6 | EGG1 | 2^8 | 7.67 |
| A/mallard/Maryland/09OS1359/2009 | H4N6 | EGG1 | 2^7 | 6.5 |
| A/mallard/Maryland/1241/2005 | H4N6 | EGG1 | 2^8 | 7.67 |
| A/mallard/Maryland/11OS952/2011 | H4N6 | EGG1 | 2^8 | 8.5 |
| A/northernshoveler/Wisconsin/11OS4110/2011 | H4N6 | EGG1 | 2^8 | 8 |
| A/mallard/Maryland/14OS1321/2014 | H4N6 | EGG1 | 2^8 | 7.5 |
| A/mallard/Wisconsin/08OS2261/2008 | H4N6 | EGG1 | 2^8 | 8 |
| A/mallard/Maryland/09OS1361/2009 | H4N6 | EGG1 | 2^8 | 8 |
| A/blue-wingedteal/Wisconsin/2741/2009 | H4N6 | EGG1 | 2^9 | 8 |
| A/green-wingedteal/Ohio/15OS5497/2015 | H4N6 | EGG1 | 2^9 | 8 |
| A/Americangreen-wingedteal/Wisconsin/11OS3592/2011 | H4N6 | EGG1 | 2^6 | 8 |
| A/Americangreen-wingedteal/Wisconsin/12OS2997/2012 | H4N6 | EGG1 | 2^9 | 8 |
| A/blue-wingedteal/Missouri/15OS5134/2015 | H4N6 | EGG1 | 2^9 | 5.5 |
| A/environment/Maryland/1101/2006 | H4N6 | EGG1 | 2^6 | 6.5 |
| A/blue-wingedteal/Missouri/15OS4803/2015 | H4N6 | EGG1 | 2^8 | 7.33 |
| A/blue-wingedteal/Illinois/10OS1562/2010 | H4N6 | EGG1 | 2^10 | 9 |
| A/mallard/Maryland/09OS1355/2009 | H4N6 | EGG1 | 2^10 | 8.5 |
| A/gadwall/Wisconsin/11OS3415/2011 | H4N6 | EGG1 | 2^10 | 8 |
| A/blue-wingedteal/Ohio/15OS5174/2015 | H4N6 | EGG1 | 2^10 | 8.67 |
| A/environment/Maryland/09OS1303/2009 | H4N6 | EGG1 | 2^9 | 8 |
| A/environment/Maryland/09OS1308/2009 | H4N6 | EGG1 | 2^9 | 8.33 |
| A/blue-wingedteal/Wisconsin/11OS2657/2011 | H4N6 | EGG1 | 2^8 | 7.67 |
| A/green-Wingedteal/Ohio/15OS4704/2015 | H4N6 | EGG1 | 2^9 | 8.5 |
| A/environment/Maryland/14OS1486/2014 | H4N6 | EGG1 | 2^9 | 8 |
| A/blue-wingedteal/Missouri/15OS4830/2015 | H4N6 | EGG1 | 2^6 | 7 |
| A/mallard/Ohio/15OS4308/2015 | H4N6 | EGG1 | 2^10 | 8.5 |
| A/blue-wingedteal/Ohio/13OS2016/2013 | H4N6 | EGG1 | 2^8 | 8 |
| A/environment/Maryland/14OS1364/2014 | H4N6 | EGG1 | 2^8 | 7 |
| A/mallard/Ohio/655/2002 | H4N6 | EGG1 | 2^9 | 8 |
| A/mallard/Maryland/06OS1110/2006 | H4N6 | EGG1 | 2^9 | 6.5 |
| A/blue-wingedteal/Iowa/13OS2338/2013 | H4N6 | EGG1 | 2^9 | 8 |
| A/mallard/Ohio/12OS5547/2012 | H4N6 | EGG1 | 2<sup>9</sup> | 8 |
| A/environment/Maryland/09OS1295/2009 | H4N6 | EGG1 | 2<sup>7</sup> | 6.67 |
| A/blue-wingedteal/Missouri/15OS4786/2015 | H4N6 | EGG1 | 2<sup>9</sup> | 8 |
| A/mallard/Ohio/15OS4302/2015 | H4N6 | EGG1 | 2<sup>9</sup> | 8 |
| A/Americangreen-wingedteal/Missouri/15OS6530/2015 | H4N6 | EGG1 | 2<sup>9</sup> | 6.67 |
| A/bluewingedteal/Missouri/14OS1868/2014 | H4N6 | EGG1 | 2<sup>9</sup> | 7.67 |

**Table S2.** Viral titers for the nasal wash samples from feral swine. The viral titers were determined by qRT-PCR.

| Virus | Pen # | Treatment | Pig ID | Necropsy | Viral shedding in nasal wash samples (Mean ± SD Log<sub>10</sub> Copies/mL) | | | | | |
| --- | --- | --- | --- | --- | --- | --- | --- | --- | --- | --- |
|  |  |  |  |  | 0 dpi<sup>a</sup> | 2 dpi | 4 dpi | 6 dpi | 8 dpi | 10 dpi |
| MO/15 | 3 | inoculated | 143 | 5 dpi | NA<sup>b</sup> | 4.68±0.38 | 3.28±0.37 | —<sup>c</sup> | — | — |
| MO/15 | 3 | contact | 150 | 21 dpi | NA | NA | NA | NA | NA | NA |
| MO/15 | 3 | inoculated | 163 | 21 dpi | NA | 5.09±0.37 | 6.28±0.47 | 3.61±0.37 | NA | NA |
| MO/15 | 5 | contact | 142 | 21 dpi | NA | NA | NA | NA | NA | NA |
| MO/15 | 5 | inoculated | 144 | 7 dpi | NA | 4.90±0.38 | 5.55±0.42 | 3.23±0.35 | — | — |
| MO/15 | 7 | contact | 147 | 21 dpi | NA | NA | NA | 4.74±0.35 | 5.39±0.29 | NA |
| MO/15 | 7 | inoculated | 148 | 21 dpi | NA | 4.70±0.42 | 4.60±0.36 | NA | NA | NA |
| OS2244 | A | inoculated | 155 | 5 dpi | NA | NA | NA | — | — | — |
| OS2244 | A | contact | 140 | 21 dpi | NA | NA | NA | NA | NA | NA |
| OS2244 | A | inoculated | 162 | 21 dpi | NA | NA | NA | NA | NA | NA |
| OS2244 | C | contact | 153 | 21 dpi | NA | NA | NA | NA | NA | NA |
| OS2244 | C | inoculated | 156 | 7 dpi | NA | 4.56±0.26 | 3.52±0.31 | NA | — | — |
| OS2244 | B | contact | 149 | 21 dpi | NA | NA | NA | NA | NA | NA |
| OS2244 | B | inoculated | 151 | 21 dpi | NA | NA | 3.38±0.97 | NA | NA | NA |
| OS5426 | C | inoculated | 159 | 5 dpi | NA | NA | NA | — | — | — |
| OS5426 | C | contact | 141 | 21 dpi | NA | NA | NA | NA | NA | NA |
| OS5426 | C | inoculated | 139 | 21 dpi | NA | NA | NA | NA | NA | NA |
| OS5426 | A | contact | 146 | 7 dpi | NA | NA | NA | NA | — | — |
| OS5426 | A | inoculated | 145 | 21 dpi | NA | NA | NA | NA | NA | NA |
| OS5426 | B | contact | 154 | 21 dpi | NA | NA | NA | NA | NA | NA |
| OS5426 | B | inoculated | 152 | 21 dpi | NA | NA | NA | NA | NA | NA |
| PBS | 1 | control | 160 | 5 dpi | NA | NA | NA | — | — | — |
| PBS | 3 | control | 164 | 5 dpi | NA | NA | NA | — | — | — |
| PBS | 1 | control | 158 | 7 dpi | NA | NA | NA | NA | — | — |
| PBS | 3 | control | 165 | 7 dpi | NA | NA | NA | NA | — | — |
| PBS | 1 | control | 164 | 21 dpi | NA | NA | NA | NA | NA | NA |
| PBS | 3 | control | 161 | 21 dpi | NA | NA | NA | NA | NA | NA |

<sup>a</sup>Days post inoculation (dpi); <sup>b</sup>Not detected (NA); <sup>c</sup>Sample's not available.

**Table S3.** Viral titers for the nasal wash samples from feral swine. The viral titers were determined by ddPCR.

| Virus | Pen # | Treatment | Pig ID | Necropsy | Viral shedding in nasal wash samples (Mean ± SD Log<sub>10</sub> Copies/mL) | | | | | |
| --- | --- | --- | --- | --- | --- | --- | --- | --- | --- | --- |
|  |  |  |  |  | 0 dpi<sup>a</sup> | 2 dpi | 4 dpi | 6 dpi | 8 dpi | 10 dpi |
| MO/15 | 3 | inoculated | 143 | 5 dpi | NA<sup>b</sup> | 4.49 | 2.79 | —<sup>c</sup> | — | — |
| MO/15 | 3 | contact | 150 | 21 dpi | NA | NA | NA | NA | NA | NA |
| MO/15 | 3 | inoculated | 163 | 21 dpi | NA | 4.75 | 5.89 | 3.19 | NA | NA |
| MO/15 | 5 | contact | 142 | 21 dpi | NA | NA | NA | NA | NA | NA |
| MO/15 | 5 | inoculated | 144 | 7 dpi | NA | 4.72 | 5.00 | 3.00 | — | — |
| MO/15 | 7 | contact | 147 | 21 dpi | NA | NA | NA | 4.40 | 5.06 | NA |
| MO/15 | 7 | inoculated | 148 | 21 dpi | NA | 4.33 | 4.24 | NA | NA | NA |
| OS2244 | A | inoculated | 155 | 5 dpi | NA | NA | NA | — | — | — |
| OS2244 | A | contact | 140 | 21 dpi | NA | NA | NA | NA | NA | NA |
| OS2244 | A | inoculated | 162 | 21 dpi | NA | NA | NA | NA | NA | NA |
| OS2244 | C | contact | 153 | 21 dpi | NA | NA | NA | NA | NA | NA |
| OS2244 | C | inoculated | 156 | 7 dpi | NA | 4.62 | 3.13 | NA | — | — |
| OS2244 | B | contact | 149 | 21 dpi | NA | NA | NA | NA | NA | NA |
| OS2244 | B | Inoculated | 151 | 21 dpi | NA | NA | 2.57 | NA | NA | NA |
| OS5426 | C | Inoculated | 159 | 5 dpi | NA | NA | NA | — | — | — |
| OS5426 | C | Contact | 141 | 21 dpi | NA | NA | NA | NA | NA | NA |
| OS5426 | C | Inoculated | 139 | 21 dpi | NA | NA | NA | NA | NA | NA |
| OS5426 | A | Contact | 146 | 7 dpi | NA | NA | NA | NA | — | — |
| OS5426 | A | Inoculated | 145 | 21 dpi | NA | NA | NA | NA | NA | NA |
| OS5426 | B | Contact | 154 | 21 dpi | NA | NA | NA | NA | NA | NA |
| OS5426 | B | Inoculated | 152 | 21 dpi | NA | NA | NA | NA | NA | NA |
| PBS | 1 | Control | 160 | 5 dpi | NA | NA | NA | — | — | — |
| PBS | 3 | Control | 164 | 5 dpi | NA | NA | NA | — | — | — |
| PBS | 1 | Control | 158 | 7 dpi | NA | NA | NA | NA | — | — |
| PBS | 3 | Control | 165 | 7 dpi | NA | NA | NA | NA | — | — |
| PBS | 1 | Control | 164 | 21 dpi | NA | NA | NA | NA | NA | NA |
| PBS | 3 | Control | 161 | 21 dpi | NA | NA | NA | NA | NA | NA |

<sup>a</sup>Days post inoculation (dpi); <sup>b</sup>Not detected (NA); <sup>c</sup>Samples not available.

**Table S4.** Viral titers for the respiratory tract tissue samples from feral swine. The viral titers were determined by qRT-PCR.

| Viral loading in respiratory tract tissues (Mean ± SD Log<sub>10</sub> Copies/g) | | | | | | | | | | |
| --- | --- | --- | --- | --- | --- | --- | --- | --- | --- | --- |
| Virus | MO/15 |  | OS2244 |  | OS5426 |  | PBS |  |  |  |
| Treatment | Inoculated | inoculated | inoculated | inoculated | inoculated | contact | control |  |  |  |
| Pig ID | 143 | 144 | 155 | 156 | 159 | 146 | 160 | 164 | 158 | 165 |
| Necropsy | 5 dpi<sup>a</sup> | 7 dpi | 5 dpi | 7 dpi | 5 dpi | 7 dpi | 5 dpi | 5 dpi | 7 dpi | 7 dpi |
| RT | 4.10±0.17 | 4.51±0.36 | NA<sup>b</sup> | 3.85±0.40 | 4.86±0.06 | NA | NA | NA | NA | NA |
| MT | 4.26±0.05 | NA | NA | NA | 6.63±0.02 | NA | NA | NA | NA | NA |
| ET | NA | NA | 3.77±0.04 | 4.18±0.20 | 6.64±0.01 | NA | NA | NA | NA | NA |
| SP | NA | NA | NA | NA | 4.84±0.01 | NA | NA | NA | NA | NA |
| TR-U | 4.74±0.05 | NA | 3.83±0.13 | NA | 3.91±0.29 | NA | NA | NA | NA | NA |
| TR-M | 4.72±0.03 | NA | 4.21±0.06 | NA | 4.25±0.02 | NA | NA | NA | NA | NA |
| TR-D | 4.67±0.03 | NA | 4.60±0.10 | NA | NA | NA | NA | NA | NA | NA |
| BR | 5.11±0.06 | NA | 7.59±0.04 | NA | 4.24±0.04 | NA | NA | NA | NA | NA |
| LCR | NA | NA | 5.98±0.05 | 3.90±0.63 | 4.43±0.04 | NA | NA | NA | NA | NA |
| LCD | NA | NA | 4.30±0.09 | 3.81±0.11 | 3.68±0.26 | NA | NA | NA | NA | NA |
| RCR | NA | NA | NA | NA | NA | NA | NA | NA | NA | NA |
| RMD | NA | NA | NA | NA | NA | NA | NA | NA | NA | NA |
| RCD | NA | NA | 5.18±0.02 | NA | NA | NA | NA | NA | NA | NA |
| RA | NA | NA | 5.70±0.02 | NA | NA | NA | NA | NA | NA | NA |

<sup>a</sup>Days post inoculation (dpi); <sup>b</sup>Not detected (NA).

**Table S5.**
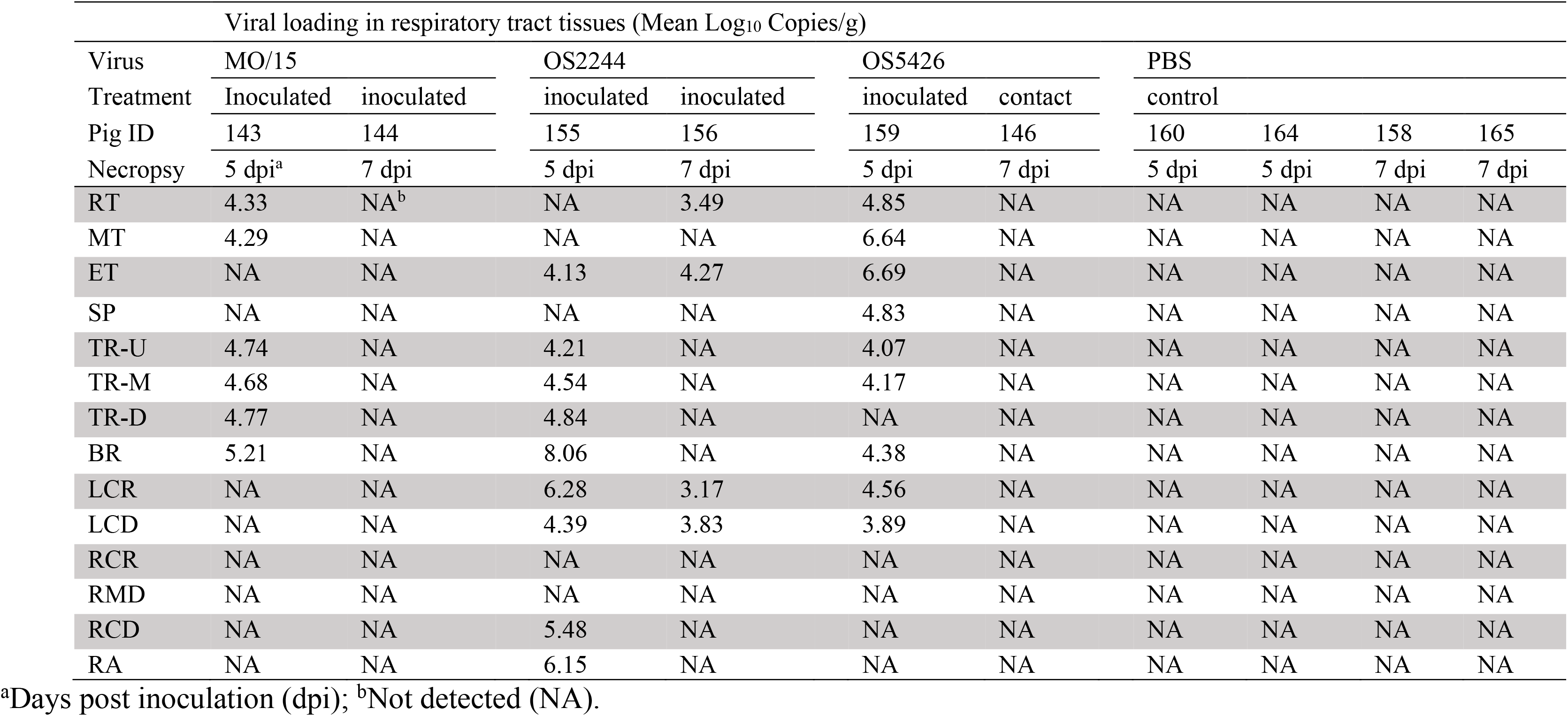
Viral titers for the respiratory tract tissue samples from feral swine. The viral titers were determined by ddPCR.

**Table S6.** The HI titers for sera collected from feral swine. The HI titer was determined against MO/15.

| Virus | Pen # | Treatment | Pig ID | Necropsy | HI titer of serum samples against MO/15 |  |  |  |  |
| --- | --- | --- | --- | --- | --- | --- | --- | --- | --- |
|  |  |  |  |  | 0 dpi<sup>a</sup> | 6 dpi | 10 dpi | 14 dpi | 21 dpi |
| MO/15 | 3 | inoculated | 143 | 5 dpi | <1:10 | —<sup>b</sup> | — | — | — |
| MO/15 | 3 | contact | 150 | 21 dpi | <1:10 | <1:10 | <1:10 | <1:10 | <1:10 |
| MO/15 | 3 | inoculated | 163 | 21 dpi | <1:10 | <1:10 | 1:160 | 1:80 | 1:40 |
| MO/15 | 5 | contact | 142 | 21 dpi | <1:10 | <1:10 | <1:10 | <1:10 | <1:10 |
| MO/15 | 5 | inoculated | 144 | 7 dpi | <1:10 | <1:10 | — | — | — |
| MO/15 | 7 | contact | 147 | 21 dpi | <1:10 | <1:10 | 1:40 | 1:640 | 1:160 |
| MO/15 | 7 | inoculated | 148 | 21 dpi | <1:10 | <1:10 | 1:160 | 1:160 | 1:80 |
| OS2244 | A | inoculated | 155 | 5 dpi | <1:10 | — | — | — | — |
| OS2244 | A | contact | 140 | 21 dpi | <1:10 | <1:10 | <1:10 | <1:10 | <1:10 |
| OS2244 | A | inoculated | 162<sup>c</sup> | 21 dpi | <1:10 | <1:10 | 1:80 | — | — |
| OS2244 | C | contact | 153 | 21 dpi | <1:10 | <1:10 | <1:10 | <1:10 | <1:10 |
| OS2244 | C | inoculated | 156 | 7 dpi | <1:10 | <1:10 | — | — | — |
| OS2244 | B | contact | 149 | 21 dpi | <1:10 | <1:10 | <1:10 | <1:10 | <1:10 |
| OS2244 | B | inoculated | 151 | 21 dpi | <1:10 | <1:10 | 1:320 | 1:320 | 1:160 |
| OS5426 | C | inoculated | 159 | 5 dpi | <1:10 | — | — | — | — |
| OS5426 | C | contact | 141 | 21 dpi | <1:10 | <1:10 | <1:10 | <1:10 | <1:10 |
| OS5426 | C | inoculated | 139 | 21 dpi | <1:10 | <1:10 | <1:10 | 1:80 | 1:40 |
| OS5426 | A | contact | 146 | 7 dpi | <1:10 | <1:10 | — | — | — |
| OS5426 | A | inoculated | 145 | 21 dpi | <1:10 | <1:10 | 1:10 | 1:40 | 1:20 |
| OS5426 | B | contact | 154 | 21 dpi | <1:10 | <1:10 | <1:10 | <1:10 | <1:10 |
| OS5426 | B | inoculated | 152 | 21 dpi | <1:10 | <1:10 | 1:80 | 1:40 | 1:20 |
| PBS | 1 | control | 160 | 5 dpi | <1:10 | — | — | — | — |
| PBS | 3 | control | 164 | 5 dpi | <1:10 | — | — | — | — |
| PBS | 1 | control | 158 | 7 dpi | <1:10 | <1:10 | — | — | — |
| PBS | 3 | control | 165 | 7 dpi | <1:10 | <1:10 | — | — | — |
| PBS | 1 | control | 164 | 21 dpi | <1:10 | <1:10 | <1:10 | <1:10 | <1:10 |
| PBS | 3 | control | 161 | 21 dpi | <1:10 | <1:10 | <1:10 | <1:10 | <1:10 |

<sup>a</sup>Days post inoculation (dpi); <sup>b</sup>Not available (—); <sup>c</sup>The pig was euthanized on 12 dpi due to leg injury.

### Supplementary Figures

**Supplementary Figure 1.**
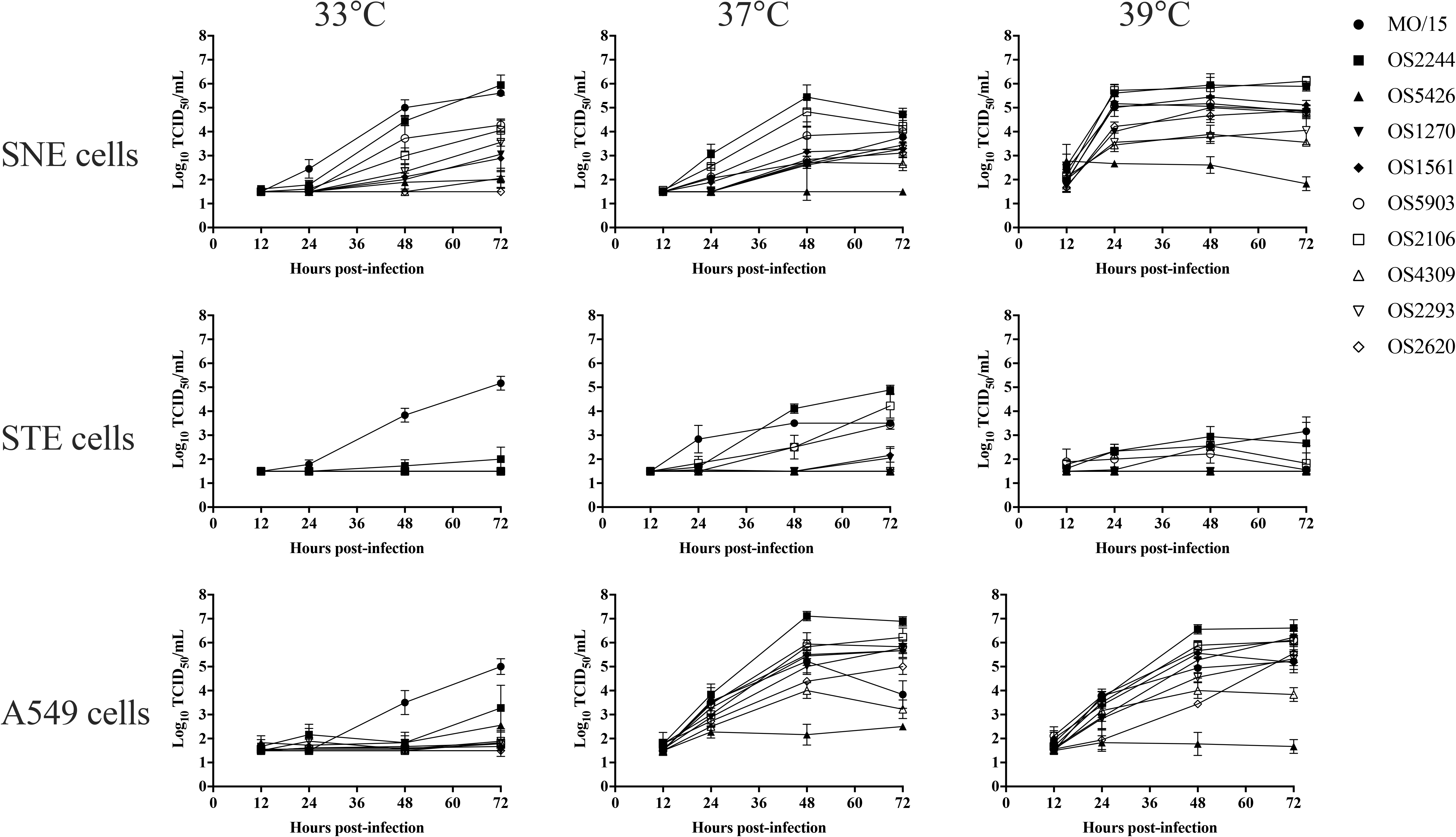
Growth kinetics of nine representative H4N6 avian isolates and a H4N6 swine isolate, MO/15, on SNE, STE, and A549 cells and at three different temperatures. Cells were infected at a multiplicity of infection of 0.001 TCID_50_/cell with the indicated viruses. Infected cells were incubated at 33°C, 37°C, or 39°C. Growth curves were determined by using the viral titers in the supernatants of infected cells obtained at 12, 24, 48, and 72 h post-inoculation. Data shown represent the mean titers ± standard errors (n = 3 cultures). Abbreviations: A/swine/Missouri/A01727926/2015 (H4N6), MO/15; A/blue-winged teal/Ohio/12OS2244/2012(H4N6), OS2244; A/blue-winged teal/Ohio/15OS5426/2015 (H4N6), OS5426; A/Mallard/Ohio/08OS1270/2008 (H4N6), OS1270; A/Blue winged teal/Illinois/10OS1561/2010 (H4N6), OS1561; A/American green winged teal/Mississippi/11OS5903/2011 (H4N6), OS5903; A/Blue winged teal/Ohio/12OS2106/2012 (H4N6), OS2106; A/Mallard/Wisconsin/11OS4309/2011 (H4N6), OS4309; /Blue winged teal/Ohio/12OS2293/2012, OS2293; and A/Blue winged teal/Missouri/11OS2620/2011 (H4N6), OS2620.

**Supplementary Figure 2.**
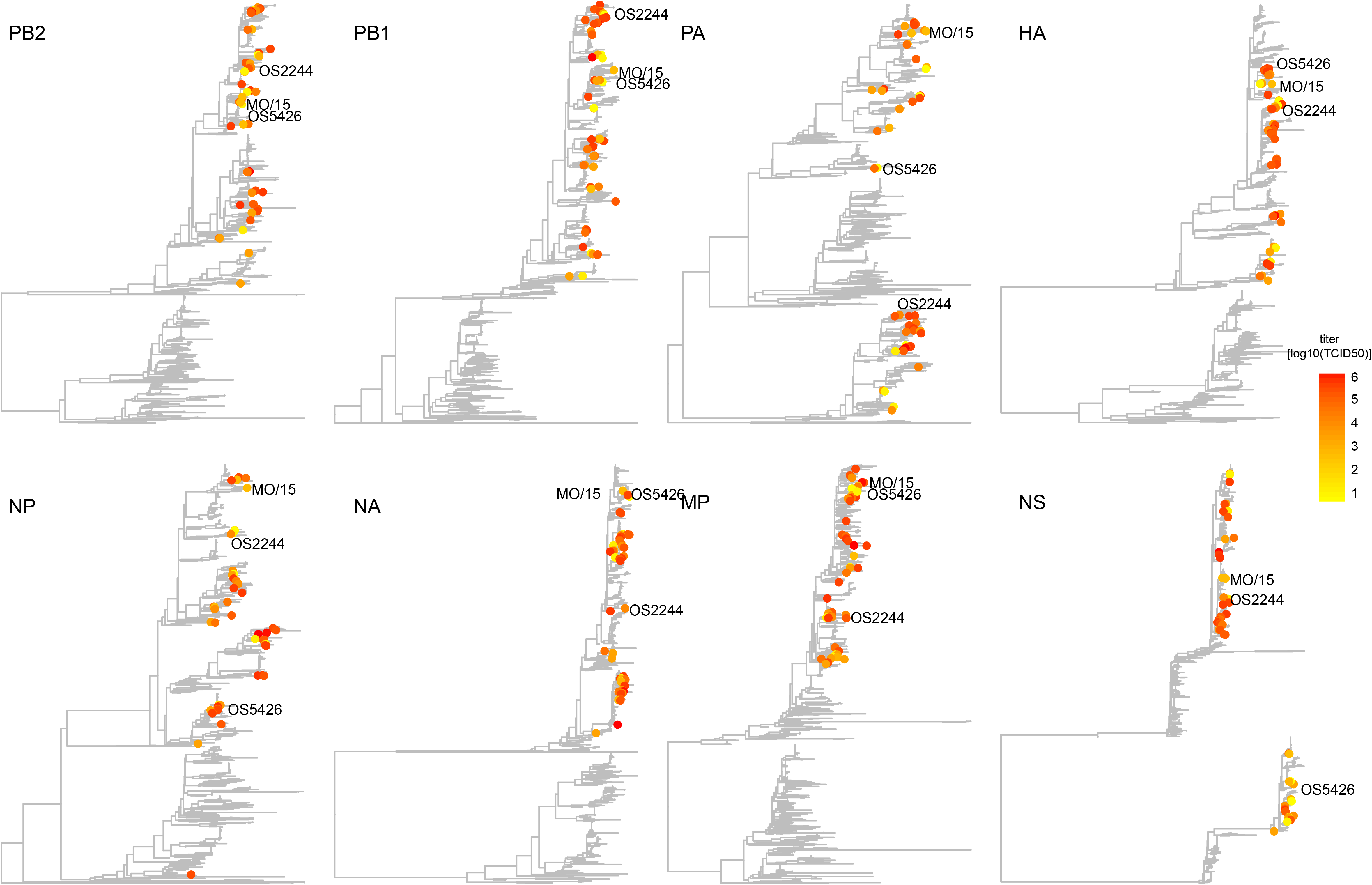
Distribution of growth phenotypes on STE cells (72 h and 37 °C) variants in phylogenetic trees. The phylogenetic trees for each gene segment was inferred using a maximum-likelihood method implemented in RAxML v8.2.9. A GAMMA model of rate heterogeneity and a generalized time-reversible (GTR) substitution model were applied in the analysis. Phylogenetic trees were then visualized by ggtree v1.6.11. Three viruses, MO/15, OS2244, and OS5425, used in animal study are annotated. Abbreviations are described in the legend of SFig. 1.

**Supplementary Figure 3.**
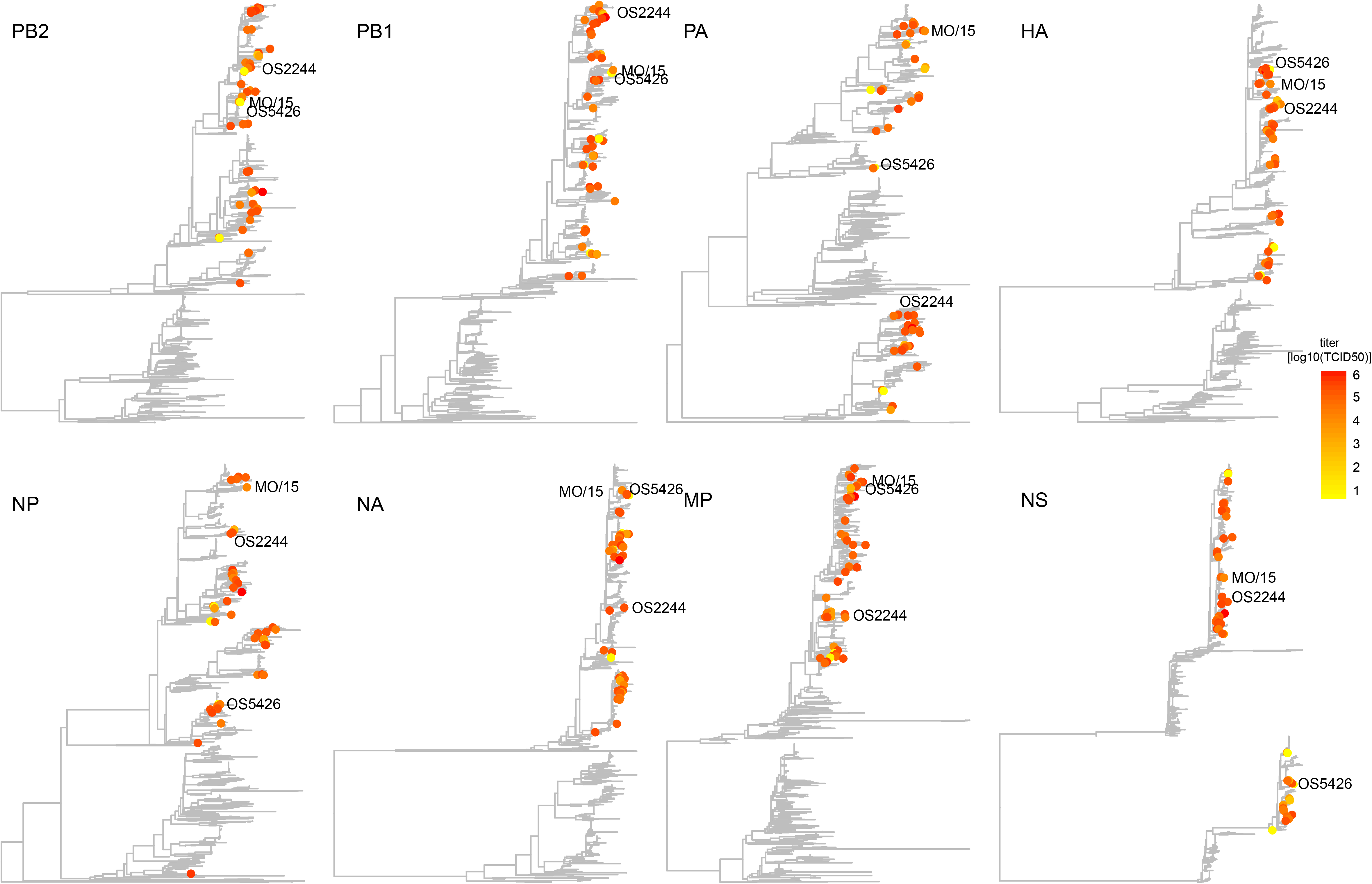
Distribution of growth phenotypes on human A549 cells (72 h and 39 °C) variants in phylogenetic trees. The phylogenetic trees for each gene segment was inferred using a maximum-likelihood method implemented in RAxML v8.2.9. A GAMMA model of rate heterogeneity and a generalized time-reversible (GTR) substitution model were applied in the analysis. Phylogenetic trees were then visualized by ggtree v1.6.11. Three viruses, MO/15, OS2244, and OS5425, used in animal study are annotated. Abbreviations are described in the legend of SFig. 1.

